# The songbird basal ganglia connectome

**DOI:** 10.1101/2025.10.25.684569

**Authors:** A. Rother, M. Januszewski, V. Jain, M.S. Fee, J. Kornfeld

**Affiliations:** Max Planck Institute for Biological Intelligence, Martinsried, Germany; Google Research, Zurich, Switzerland; Google Research, Mountain View, CA, USA; Department of Brain and Cognitive Sciences, McGovern Institute for Brain Research, Massachusetts Institute of Technology, Cambridge, MA, USA; MRC Laboratory of Molecular Biology, Francis Crick Avenue, Cambridge Biomedical Campus, Cambridge CB2 0QH, UK; Max Planck Institute for Medical Research, Heidelberg, Germany

## Abstract

The basal ganglia (BG) play an essential role in shaping vertebrate behavior, ranging from motor learning to emotions, but comprehensive maps of their canonical synaptic architecture are missing. In mammals, three main neuronal pathways through the BG have been described - the direct, indirect, and hyperdirect pathways - which together orchestrate many aspects of learning and behavior. In songbirds, cell types associated with striatal and pallidal components appear intermingled in a single basal ganglia nucleus, Area X, essential for song learning. This allows for the dense reconstruction of the entire circuit within a compact volume. Here, we introduce the first vertebrate basal ganglia connectome, comprising over 8,500 automated neuron reconstructions connected by about 20 million synapses. High image quality and automated reconstruction allowed analysis with minimal manual proofreading. Based on direct anatomical measurement of synaptic connectivity, we confirm that a direct, indirect and hyperdirect pathway can be traced through Area X. However, detailed morphological and connectomic analysis revealed no clearly distinct direct and indirect medium spiny neuron subpopulations, and a dominance of the direct and hyperdirect pathway. In addition to previously identified neuron types in Area X, we could distinguish three novel GABAergic neuron types, two of which are major output targets of GPe neurons, leading to novel feedback circuitry within Area X. We further found unexpectedly strong neuronal interconnectivity and recurrency between neurons associated with all pathways. Our data thus challenge the universality of the view of the basal ganglia as an information processor organized into discrete feedforward pathways.

## Main text

While BG circuitry are especially important for motor control, including learning^1–6^, they appear to be involved in most brain functions, including emotional processing^7^. Not surprisingly, a large spectrum of devastating neurological disorders ranging from Parkinson’s disease^8^ to addiction^9^ has been directly linked to BG dysfunction.

From a circuit perspective, the BG consist of a canonical repeating motif, similar to the cortical and thalamic circuits to which they are anatomically and functionally linked^4,10^. Historically, this canonical motif includes the direct, indirect and hyperdirect BG pathways, which in the mammal span the striatum, globus pallidus internal (GPi) / substantia nigra pars reticulata and external (GPe), and the subthalamic nucleus (STN). These regions are, at least in mammals, macro-anatomically separated (Fig.1a)^11^.

The direct and indirect pathways start with cortical axons innervating striatal medium spiny neurons (MSNs), that in turn innervate and converge on the pallidal GPi output layer (Fig. 1b)^12,13^. In mammals, MSNs can be separated into direct and indirect pathways, which appear to largely express either D1 or D2 dopamine receptors^14–16^. In the direct pathway, MSN activation ultimately disinhibits the thalamus by inhibiting the GPi/SNr neurons^17^, which, broadly speaking, promotes behavior. In contrast, indirect pathway MSNs inhibit GPe neurons, increasing GPi neuron activity, resulting in inhibition of the thalamus. Thus, the indirect pathway has been seen playing an opposing role to the direct pathway^18^. However, there are complexities to this simple story, revealed by parallel pathway activation^19^; and even anatomically, direct pathway MSN were observed to send axonal arborizations to GPe^20^.

**Figure 1.**
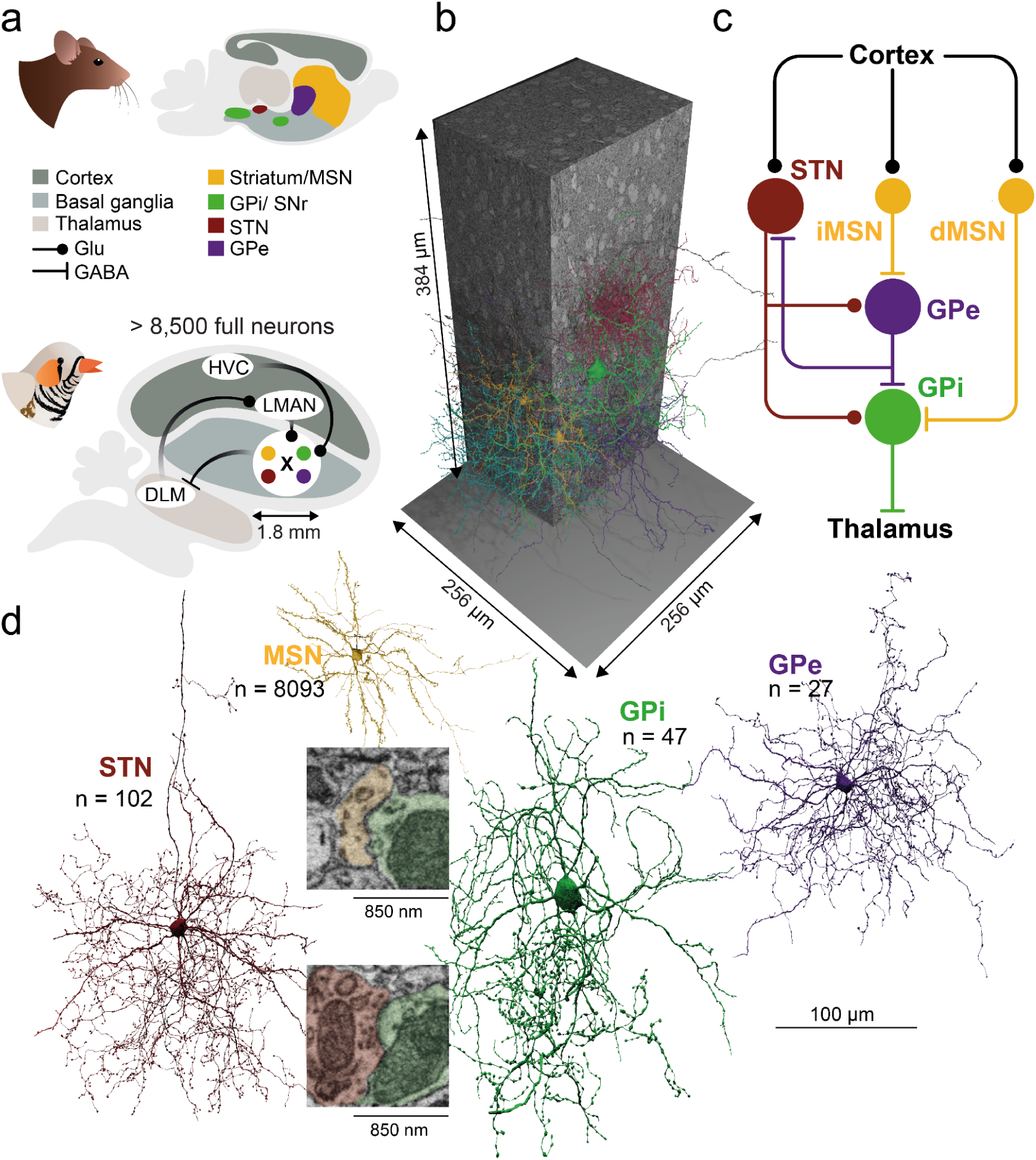
The basal ganglia in mammals and songbirds. **a** Macro-anatomical organization of the mammalian BG. **b** Canonically described main cellular BG pathways in mouse. Inset raw data coordinates: MSN-GPi:19704, 14822, 7295; STN-GPi: 17598, 12925, 5037 **c** Left: The zebra finch song system, organized in discrete nuclei, with its main BG component Area X. BG cell types are intermingled in Area X. Right: Dimensions of the connectomic Area X dataset. **d** Fully automatic reconstructions of the BG main cell types identified in this study using flood-filling networks ^41^.

In addition to the direct and indirect pathways, there is a hyperdirect pathway in mammals that does not involve MSNs, but provides an excitatory effect of cortical inputs onto GPi neurons through intervening excitatory neurons of the subthalamic nucleus (STN)^13,21^. The net effect of this pathway may be to inhibit pallidal target regions, potentially arresting ongoing behaviors, although some debate over its specific function persists^22,23^.

Previous studies have utilized a number of tracing methods to elucidate the BG circuits^24–27^. To fully reveal the circuit structure of the basal ganglia, a dense connectomic characterization, currently lacking across all vertebrate species, is needed.

The zebra finch is an important model organism for the study of motor learning and performance, due to the well delineated brain structures underlying its stereotyped but learned courtship song^28–32^. The singing behavior is coordinated by brain nuclei that include cortical/pallial and subcortical structures^33^. As part of the anterior forebrain pathway, the cortical nuclei of the lateral magnocellular nucleus of the anterior nidopallium (LMAN) and HVC (used as proper name), project to the main basal ganglia component, Area X which in turn projects to the median portion of dorsolateral thalamus (DLM). DLM then projects back to LMAN, forming a classic cortico-basal ganglia-thalamocortical loop.

The macro-anatomical separation of BG cell types into striatal and pallidal areas is not universal across vertebrates. Multiple cell types, GPe, GPi, STN and MSNs associated with the spatially separated BG areas in the mammal were identified electrophysiologically and morphologically inside of Area X (Fig. 1c)^34–39^. Since MSN, GPe, and STN refer to projection neurons in mammals, they will not be referred to as interneurons in this paper, even though they do not project outside of Area X. Additionally, single-cell fills revealed a class of striatal fast spiking (FS) neurons, as well as low threshold spiking (LTS), and cholinergic, tonically active neurons (TANs)^34^. Each of these has been tentatively identified in electrophysiological recordings in behaving birds^36^. Single nucleus sequencing revealed eight clusters of GABAergic neurons, a large population of MSNs co-expressing D1 and D2 receptors, plus a cholinergic TAN^40^. Apart from identifying these cell types in Area X, little is known about their connectivity. This raises an intriguing opportunity to study an entire BG circuit within a small brain volume accessible with connectomics.

We describe here a connectome from zebra finch Area X, reconstructed at synaptic resolution using automated volume EM and data analysis methods^41–44^. Our dataset includes a comprehensive annotation of cell types: putative incoming cortical axon fragments ^45^, MSNs, local inhibitory interneurons, excitatory STN-like neurons, local GPe-like neurons, as well as putative GPi-like output neurons. During cell typing, we encountered several inhibitory interneuron types with clearly distinct morphological and connectomic profiles, which had not been previously identified.

We tested the hypothesis that Area X harbors the BG pathways described for mammals. Starting from morphologically classified GPi and GPe neurons, we worked backwards to identify direct and indirect pathway MSNs, as well as connections to and from STN-like neurons. Surprisingly, MSNs do not form distinct subpopulations, as suggested by mammalian D1 and D2 MSN separation^14,46^. Furthermore, instead of an organization of synaptic connectivity in a clear feedforward fashion, we found prominent recurrent local connections, especially those involving the newly described interneuron types, supporting a novel view of BG-internal computations, likely crucial for learning and behavior^27,47^.

## Results

The EM dataset that our connectome is based on was acquired with serial block-face electron microscopy ^44^ from an adult male zebra finch (> 120 dph). The data set spans a volume of 256 x 256 x 384 µm^3^ with a voxel size of 10 x 10 x 25 nm^3^ (Fig. 1b). The dense reconstruction comprises 8,576 fully reconstructed neurons, defined as those with a soma and at least 200 µm of reconstructed axon and 200 µm of dendritic pathlength. It was based on flood-filling networks^41^, complemented by an automated annotation of synapses (Extended Data Fig. 1, Extended Data Table 1), mitochondria and vesicle clouds ^42^. Instead of expensive manual proofreading of all neurons, we performed a comprehensive estimate of remaining error rates (Extended Data Table 2, 3, Extended Data Fig. 2a). We manually inspected 775 neurons for false mergers, yielding a false merger rate of 0.0096 per mm of skeleton path length (Extended Data Table 3). The automated synapse detection was also manually validated (F1-score of 0.88 for all synapse sizes, 93% true positives for the used filtering criteria, Extended Data Fig. 1d). Cell type and compartment classifications (Extended Data Fig. 3) are based on previously described supervised classifiers based on morphology (see methods, Table 1, Extended Data Tables 4-6) ^42^. In total, the connectome comprises the following putative cell types: 8,093 MSNs, 27 GPe, 47 GPi, 102 STN, 37 LTS, 8 TAN, and 262 putative local inhibitory interneurons. It also includes several glial cell types and migratory neurons (see Methods for inclusion criteria, Fig. 1d, Fig. 2a)^48^ in preparation). Our dataset also includes more than 310,000 reconstructed axons (> 50 µm pathlength, see Extended Data Fig. 4 for fragment size distributions) classified as originating from structures outside Area X (see Methods), including HVC (n=246,153), LMAN (n=19,186) and ventral tegmental area (VTA, DA axons, n=53,522). Altogether, these components make this dataset one of the largest comprehensively annotated vertebrate connectomes to date.

**Figure 2.**
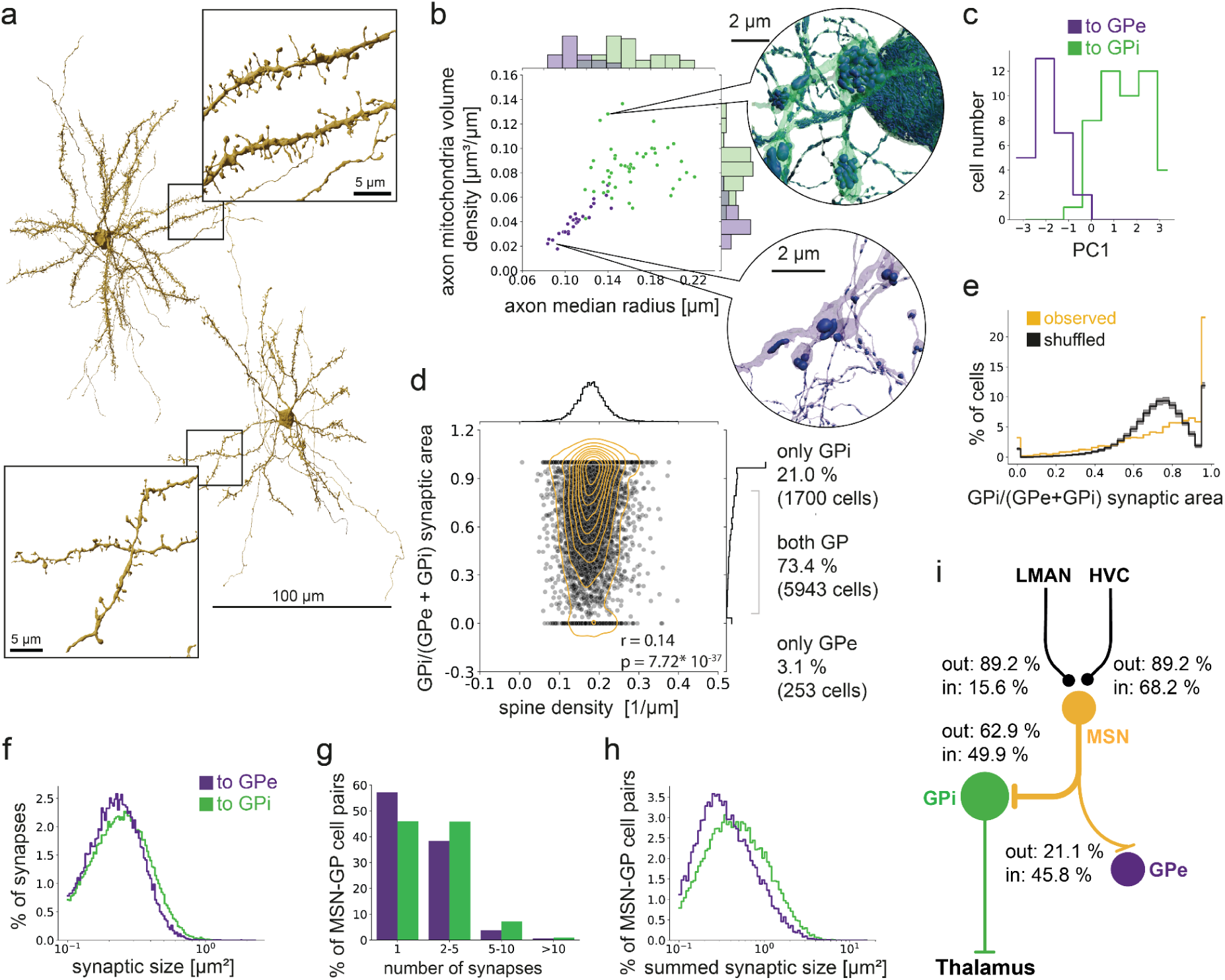
Connectomic identification of the direct and indirect pathway on the level of MSNs. **a** The densely spiny MSN cell type (see insets) can be readily identified in Area X. Link to 3D visualizations. **b** Pallidal cell types can be separated into at least two populations, consistent with electrophysiological data ^37^, putative GP and GPe based on axon mitochondria volume density, axon median radius. Insets show mitochondria in two example cells. **c** Pallidal cell types can be separated in a PCA based on parameters in b, axon myelin fraction and soma diameter (see Extended Data Fig. 5). **d** MSNs show a weak preference to connect to one pallidal cell type, and connectivity does weakly correlate with morphological features such as spine density. MSNs that did not project to either GPe or GPi were excluded (2.4%, n=197). **e** GPi preference of MSN plotted against shuffled synapses. Black curve shows mean over 100 iterations, gray indicates 95% confidence interval. **f** MSNs connect to GPi through slightly larger synapses, **g**, with more parallel synapses, and **h**, larger total synaptic area per pre-post neuronal connection. **i** Schematic representation of the direct pathway together with MSN-GPe connections with synaptic area between the shown cell types normalized to their corresponding incoming and outgoing synaptic area (see Fig. 6f, g).

**Table 1.**
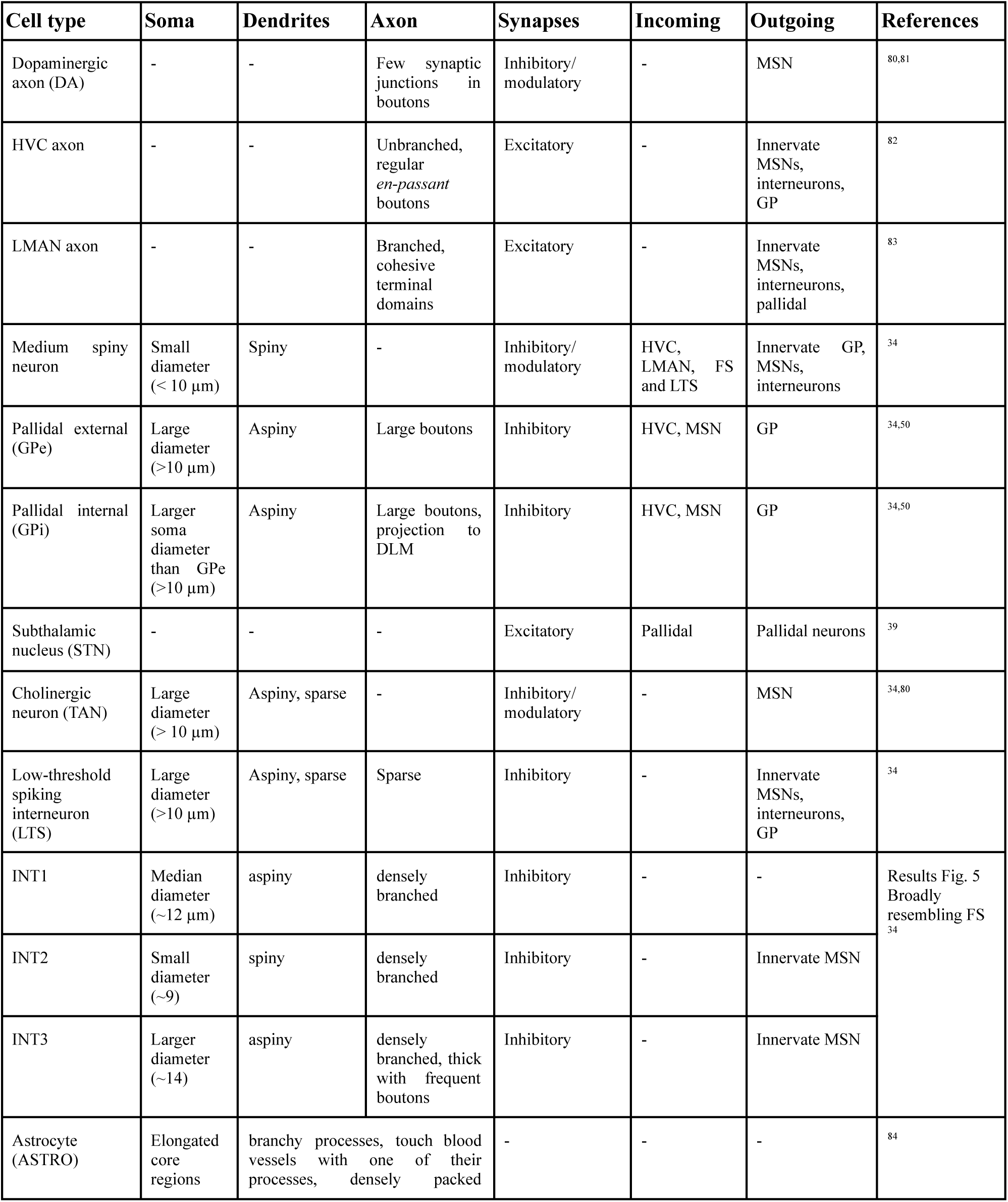
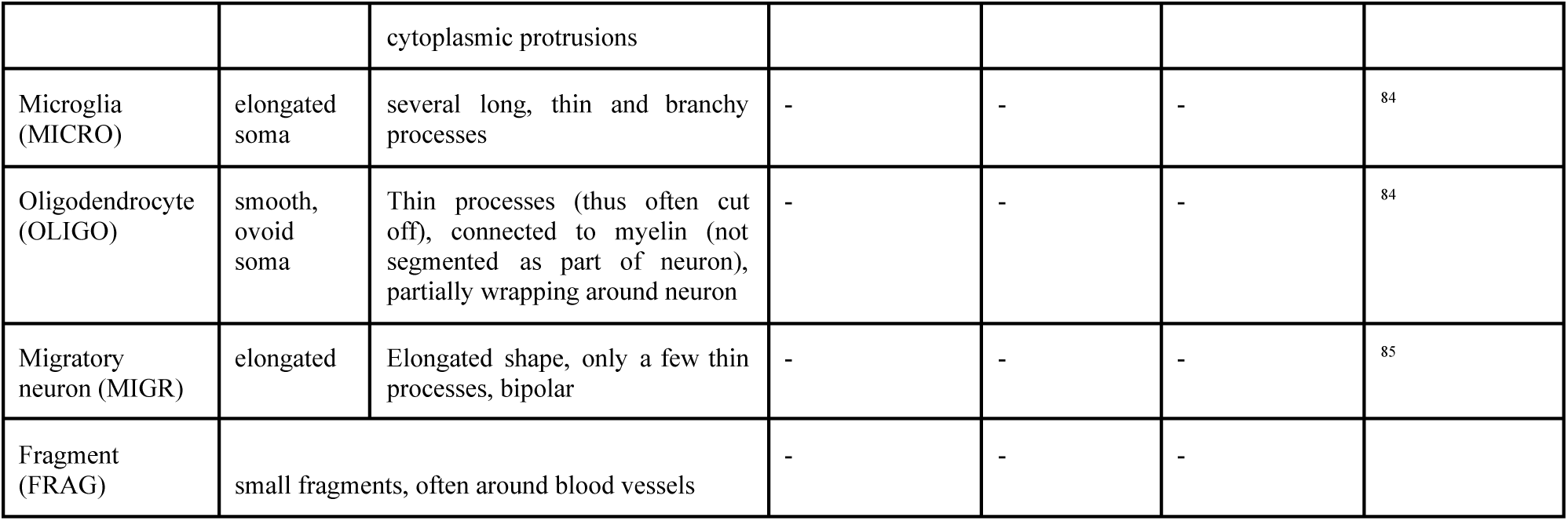
Cell types and characteristics of neurons.

The BG pathways are usually described to start with the excitatory cortical innervation of direct and indirect pathway MSNs, and indeed our data show a strong innervation of MSNs by the avian cortical analogs LMAN and HVC. Specifically, MSNs receive 15.6% of their total incoming synaptic area from LMAN, and 68.2% from HVC (see Extended Data Fig. 2c, d for an analysis of the influence of different axon fragment length filtering criteria and Extended Data Fig. 5a for synaptic area distributions for individual MSNs), and only 0.2% from local excitatory neurons (for details on HVC and LMAN connectivity to MSN, see companion paper^49^).

To analyze the distinct BG pathways downstream of these cortical inputs, we next identified GPi neurons (putative thalamus-projecting pallidal cells) and GPe neurons (putative pallidal cells that make synapses only within Area X). These two pallidal neuron types showed clear morphological differences in our dataset (Fig. 2b, c and Extended Data Fig. 5b), as previously described^50^. Both the GPe and GPi types were the primary synaptic target of MSNs (21.1% and 62.9%, respectively, of outgoing MSN synaptic area, Extended Data Fig. 5a), as expected for a canonical BG circuit. Individual MSNs made, on average, 10 synapses (9.6+-7.2) onto GPi neurons and 4 synapses onto GPe neurons (3.54+-5.4), while, on average, individual GPi neurons received over 1,000 synapses (1624+-881) from hundreds of MSNs (671+-298). Conversely, MSNs form the primary source of synaptic input to GPi and GPe neurons (45.8% for GPe, 49.9% for GPi).

We next wondered whether we could predict which MSNs project to GPi and which project to GPe, based on MSN morphology, because differences in spine density and dendritic length have been associated with direct and indirect pathway MSNs in mammals^46,51^. We found that the 8,093 reconstructed MSNs varied strongly in their morphological properties, such as spine density and dendritic length (Fig. 2a, Extended Data Fig. 5c, Extended Data Table 8). These morphological parameters showed only a weak association with GPe or GPi connectivity (r = 0.14, p-value < 0.001, Fig. 2d, Extended Data Fig. 5e). In fact, most near-completely reconstructed MSNs (see Methods, 73.4%), made synaptic contact with both pallidal types, showing substantial connectivity overlap between the direct and indirect pathway in our data (Fig. 2d)^26,52^. Only 21% of MSNs were connected exclusively to GPi and only 3% were connected exclusively to GPe. It should be noted that both these numbers are likely somewhat inflated due to incomplete reconstruction of axons at the dataset boundary (for controls see Extended Data Text, Extended Data Fig. 5-7). In total, 94.4% of MSNs project to the GPi neurons identified in our dataset, suggesting that the direct pathway predominates in the songbird Area X.

Given the large number of MSNs that project to both pallidal cell types, we wondered if individual MSNs are selective for GPi or GPe as postsynaptic partners. In fact, 35.6% of MSNs are highly selective for GPi outputs (synapse ratio to GPi >= 0.9), while this degree of selectivity would be expected in only 17.6+-0.35% of MSNs if the identity of GPi and GPe postsynaptic partners were randomly shuffled (100 iterations, Fig. 2e, see Methods). The observed data shows a peak in the GP ratio around 0.9, while in the shuffled data the peak is around 0.75, which is close to the GPi/(GPe + GPi) synaptic area fraction. Whether the identified selectivity is explained by specific molecular targeting, or the simple fact that an axon forming a synapse with a dendrite has a higher chance of connecting to that dendrite again for geometric reasons, remains to be tested.

Given the selectivity of some MSNs for targeting GPi neurons, we next analysed the distribution of synapse sizes and the number of multi-synaptic connections from MSN to GPi or GPe. We found that MSN to GPi connectivity is significantly stronger than MSN to GPe connectivity. First, MSN-GPi synapses are substantially larger, and there is a larger tendency to form multisynaptic connections with GPi neurons. This leads to MSN to GPi connections with larger total synaptic areas, in comparison to MSN to GPe (Fig. 2f-h, Wilcoxon-ranksum test, p-values < 0.001, Extended Data Table 8).

In summary, we find that MSNs receive converging cortical inputs from HVC and LMAN, and connect primarily to GPi neurons (>94%), indicating that the direct pathway is the most prominent pathway through Area X (Fig. 2i). While MSNs also connect to GPe neurons, in line with the existence of a weak indirect pathway, we find little evidence of a distinct indirect MSN population at the level of morphology or output connectivity.

We next wondered whether Area X includes an indirect pathway from GPe to GPi neurons (Fig. 3a), either through a population of STN neurons (GPe-STN-GPi), or a ‘short’ indirect pathway (GPe to GPi). We analyzed 102 putative STN neurons (1.2% of total) with a morphology resembling mammalian STNs^53–55^, previously hypothesized to exist in Area X based on electrophysiological recordings^39^. The STNs receive input from GPe neurons, however this projection represents only 6.9% of the STN incoming synaptic area and only 16.8% of the GPe outgoing synaptic area (see Fig. 3b-g for connection statistics). These observations suggest that the classical GPe-STN-GPi indirect pathway is weak in Area X. We also find that the ‘short’ indirect pathway is effectively non-existent in Area X; GPe synapses onto GPi neurons constitute only 7.5% of GPe synaptic output area and only 0.8% of GPi synaptic input area. In addition, only a small number of GPi neurons receive somatic GPe input (13/47 neurons, 28%). While GPe neurons contribute only weakly to an indirect pathway, we will show shortly that they are strongly interconnected with a novel population of local inhibitory neurons (INT2 and INT3, 67.7% of fractional synaptic input area to GPe, Fig. 5).

**Figure 3.**
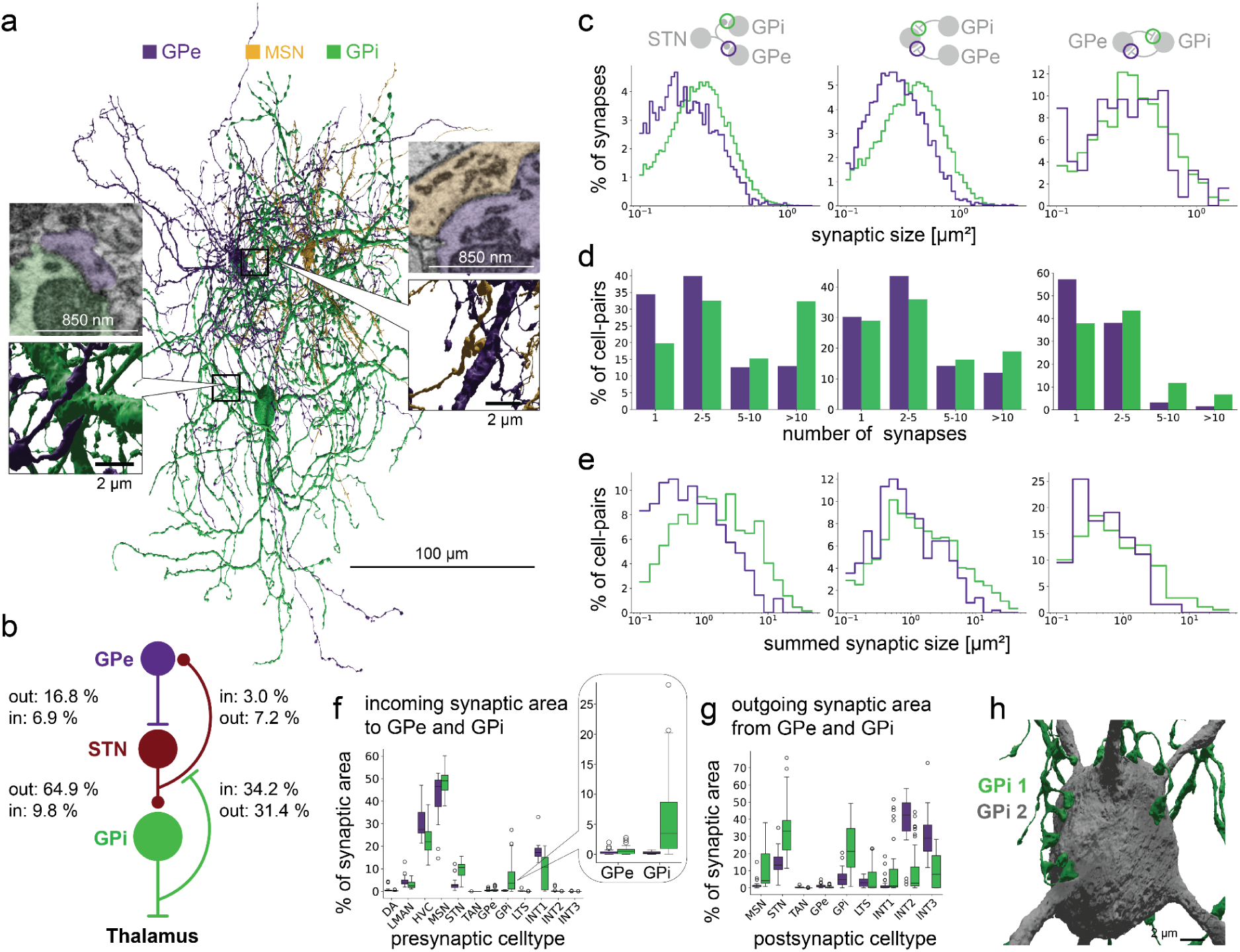
The indirect pathway in the songbird Area X. **a** Automated reconstructions of exemplary GPi and GPe neurons, and a connected MSN (3D visualizations). Insets show an MSN to GPe synapse (top:12139, 10877, 3292) and a GPe to GPi synapse (bottom: 13502, 17089, 6174). **b** Schematic representation of the indirect pathway starting at GPe with summed synaptic area in relation to corresponding incoming and outgoing synaptic area per cell type (see Fig. 6f, g), also for the connectivity from GPi to STN to GPe. **c-e** Comparison of synaptic area of individual synapses and multisynaptic connections from STN to GPe and GPi (left), STN from GPe and GPi (middle) and GPe to GPi (and vice versa, right). **f,g** Incoming and outgoing synaptic connectivity of GPe and GPi. Inset shows the incoming synaptic area to GPe and GPi. h ‘Claw’ - like, multiple soma synapses from one GPi to another (3D visualizations).

In contrast to the weak indirect pathway, we did find strong evidence for a hyperdirect pathway. In mammals, the hyperdirect pathway classically comprises cortical projections onto STNs (Fig. 4a), which excite GPi neurons, capable of putting a ‘brake’ on imminent behaviors ^55^. Correspondingly, in our dataset we find that putative STNs receive a strong projection from LMAN and HVC (16.3% and 20.7% of fractional synaptic input area to STN, Fig. 4b, c), and STNs form large excitatory synapses onto GPi neurons, representing 64.9% of total STN synaptic output (see Fig. 3b-g; STN to GPi is 9.8% of GPi input area).

**Figure 4.**
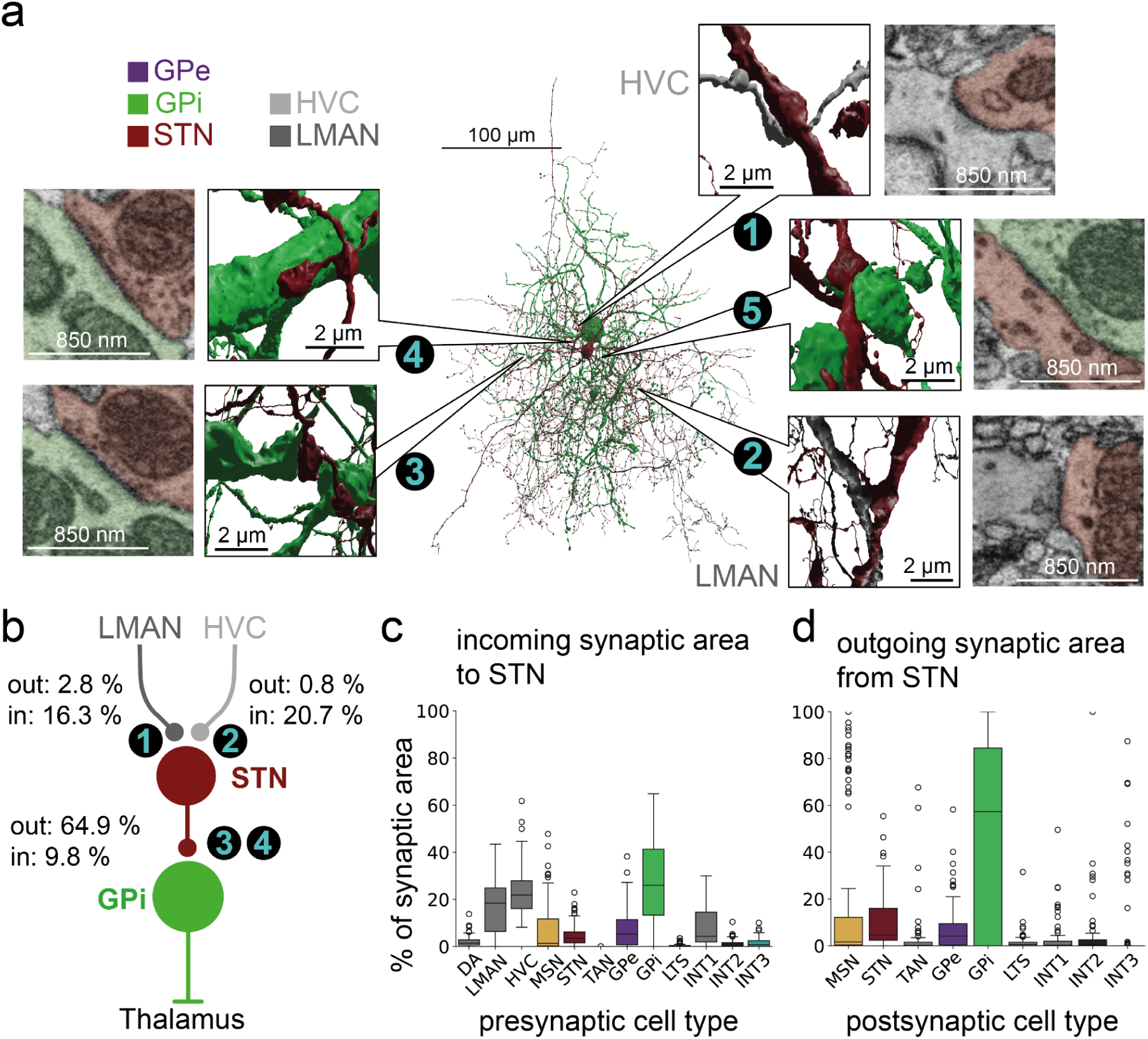
Connectomic identification of the hyperdirect pathway from cortical inputs to pallidal output neurons. **a** Reconstruction of neurons of the hyperdirect pathway (3D visualizations). Insets show asymmetric ultrastructure 1-4 ^57,58^. 1: HVC to STN synapses: 13018, 8939,7226; 2. LMAN to STN synapse: 16088, 14773, 4515; 3, 4: STN to GPi synapses: 7889, 8920, 6373; 11284, 13200, 69875; 5. GPi to STN synapse (symmetric ultrastructure): 14381, 9867, 6235 **b** Schematic representation of the hyperdirect pathway with summed synaptic area in relation to the corresponding incoming and outgoing synaptic area per cell type (see Fig. 6f, g). **c** Numbers in b,c refer to synapses shown as insets in a. **c** Cell-type level characterization of incoming and outgoing synaptic are to and from STN-like neurons.

Beyond the three canonical BG pathways described above, direct, indirect and hyperdirect pathways, we have also found evidence for a number of non-canonical features in Area X circuitry. First, we find significant direct cortical innervation of both GPi and GPe neurons, in line with reports in rat^56^. For example, both GPi and GPe neurons are also substantially innervated by HVC axons. Depending on axon fragment filtering criteria (see also Extended Data Fig. 2 and 4), these input fractions range from 28.7% to 54.7% of total synaptic input area in GPe and 20.5% to 46.1% for GPi (Extended Data Fig. 2b,c), indicating a strong cortical innervation with still unclear function. Functionally, the large HVC input to GPi and GPe neurons could account for the dramatic increase in GP firing rates during singing^37^.

A second non-canonical finding is a strong backprojection from GPi to STN, unreported in mammalian literature. This feedback comprises 31.4% of synaptic output area from GPi and 34.2% of synaptic input area to STN (Fig. 4c,d). Compared to GPe-STN and STN-GPe connections, GPi-STN and STN-GPi connections have larger synapses, and more multisynaptic connections thus leading to a larger synaptic area between individual cells (Wilcoxon ranksum test, all p < 0.01, Fig. 3 c-e, Extended Data Table 8). This recurrent synaptic coupling suggests an important functional role for the STN-GPi network.

Unexpectedly, we also identified a strong local GPi network (see Extended Data Fig. 8). The second largest output target of GPi are other GPi neurons (25.5%), which also constitute 6.9% of incoming synaptic area. These GPi to GPi connections are often multi-synaptic (see also Extended Data Fig. 8) and in part facilitated by several, ‘claw-like’ soma synapses (Fig. 3h, Extended Data Fig. 9c)^50^, which might play an important role in coordinating functional GPi output.

Our analysis of the Area X connectome revealed several additional neuron types, based on morphological and connectivity features. Consistent with mammalian and avian striatal anatomy^59,60^, we identified sparsely arborizing interneurons in line with descriptions of large tonically active cholinergic interneurons (TANs) ^34^ (Extended Data Fig. 12). These putative TANs receive inputs almost exclusively from MSNs and HVC axons (input fraction 46.3%, 36.8%, respectively). They project nearly exclusively back to MSNs (output fraction 96.2%, Extended Data Fig. 12c). While TANs show no evidence of reciprocal interactions with the same MSN cells (Extended Data Fig. 10 g, h, Extended Data Table 10), it is noteworthy that TANs receive input from a large number of MSNs (median of 236.5 MSNs), but project synaptically to only a small fraction of MSN (median of 18.5). However, we have noticed that TANs, similar to DA axons, distribute their synaptic vesicles along the entire axon, even outside of clear synaptic loci (data not shown), consistent with previous suggestions of volume transmission ^61^ by these neurons. We have additionally identified another sparsely arborizing interneuron similar to the low-threshold-spiking “LTS” type previously described ^34^ (Extended Data Fig. 13). These neurons receive most of their synaptic input from GPi neurons (input fraction 29.6%, Extended Data Fig 12c), and project mostly to MSN neurons (output fraction 68.1%).

Previous studies of songbird basal ganglia revealed the existence of fast-spiking interneurons with “FS-like” appearance, compact aspiny dendrites and densely arborizing axons with regular boutons (Fig. 5a)^34,62^. In our dataset, quantitative morphological analysis revealed three clearly separable clusters of FS-like cell types (Fig. 5b,c, Extended Data Fig. 11), with remarkably different connectomic profiles (Fig. 5d): we refer to these as INT1, INT2, and INT3. Interestingly, the INT1 subtype broadly innervates pallidal neurons and other INT1 neurons, while the other two subtypes (INT2, INT3) target almost exclusively MSNs. Most INT3 neurons (54%) exhibit autapses onto their somata or proximal dendrites, which the other two types rarely show (see Extended Data Table 11). These differences in outgoing inhibitory connectivity are accompanied by large differences on the input side—INT1 neurons receive inputs largely from LMAN axons (25.4%) and other INT1 neurons (22.8%), a significant fraction of which terminate on the large dendritic spines of INT1 neurons (Extended Data Fig. 9g). In contrast, INT2 inputs are dominated primarily by HVC (31.9%) and GPe/GPi (23.0% and 16.2%), while INT3 inputs are predominantly from HVC (59.4%). Finally, INT2 and INT3 neurons are the primary output target of GPe neurons, comprising 67.7% of total synaptic output of these pallidal neurons. These findings suggest that INT1 neurons have a distinct circuit function compared to INT2 and INT3 neurons (Fig. 6).

**Figure 5.**
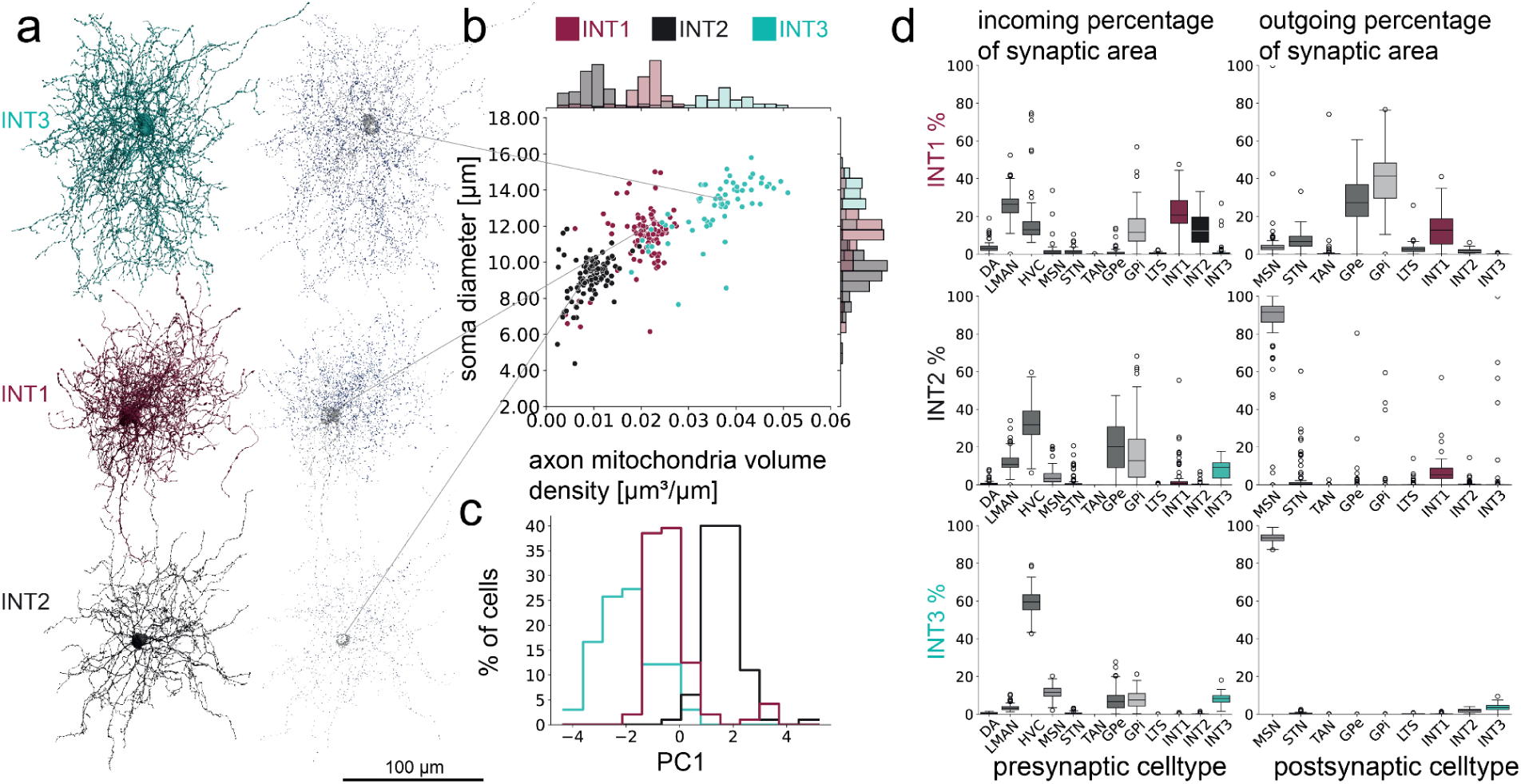
Connectomic analysis reveals diverse local interneuron types in Area X. **a** Three types of inhibitory interneurons with dense arborizations can be distinguished, all resembling broadly the fast-spiking interneuron morphology of the BG, next to visualizations of mitochondria throughout the cell (3D visualizations). Axonal mitochondria is blue, dendritic and somatic mitochondria in gray. n_INT1_ = 96, n_INT2_ = 100, n_INT3_ = 66. **b** Separation of three cell types based on soma diameter and axonal mitochondria volume in relation to axonal skeleton pathlength. **c** Separation of three cell types based on one principal component based on soma diameter, axonal mitochondria density, axon median radius and dendritic spine density (see Extended Data Fig. 11). All three interneuron types are separate from MSN (see Extended Data Fig. 6c). **d** Connectomic characterization of their synaptic inputs and output.

**Figure 6.**
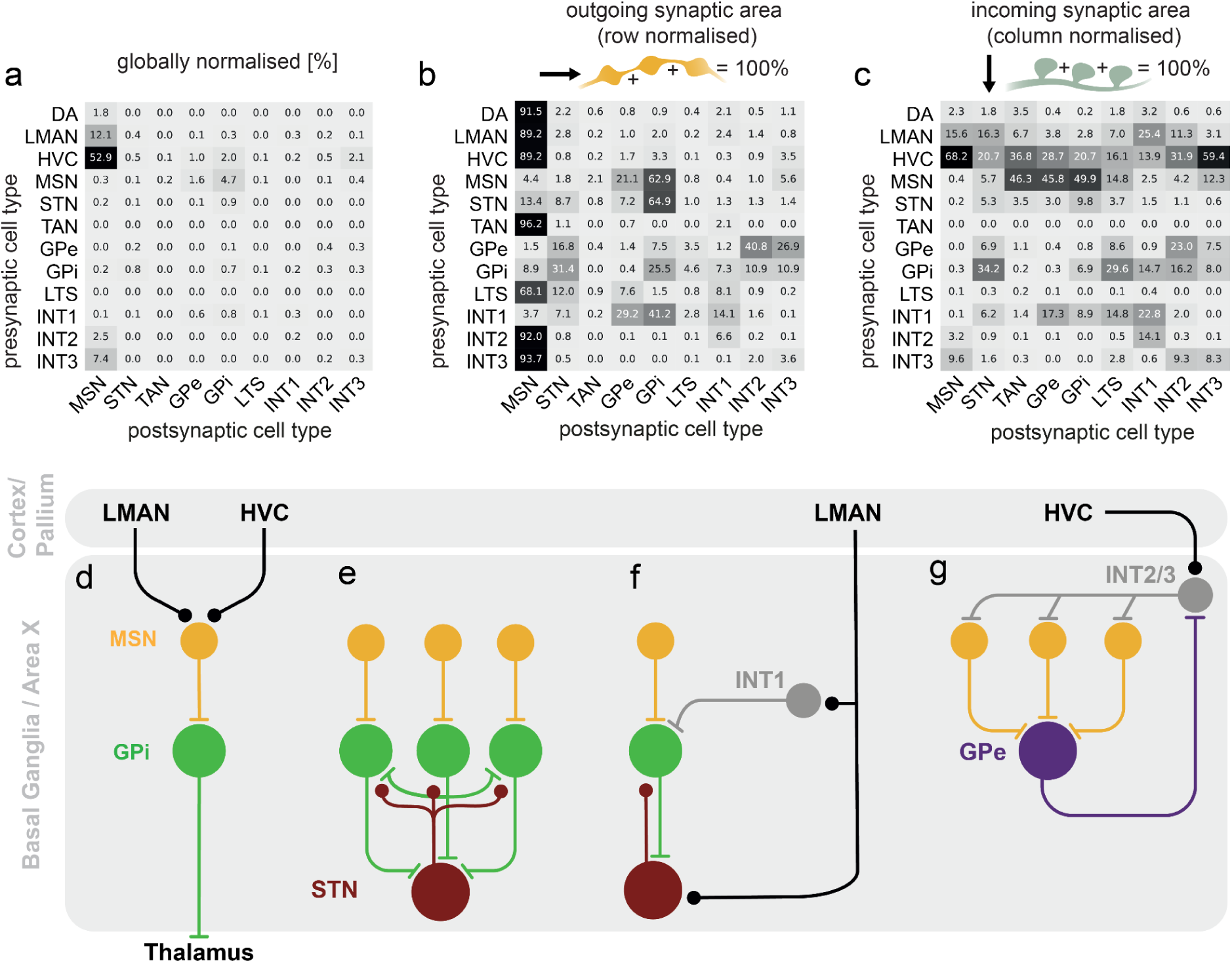
**a-c** Cell-type level connectivity matrices normalized by outgoing synaptic area (rows), incoming synaptic area (columns) and total synaptic area. **d** Schematic of the direct pathway through Area X. **e** MSNs drive a local recurrent network in which STNs and GPis interact. **f** LMAN influences GPis through inhibition via INT1s and excitation via STNs, which are part of the hyperdirect pathway. **g** Feedforward inhibition of MSNs from HVC axons through INT2/3s, and a feedback circuit from MSNs to GPes to INT2/3s cells back to MSNs.

Throughout the results so far, we have represented the pattern of connections of specific neuron types using two different views (Fig. 3f,g), a normalized distribution of synaptic output areas, and a normalized distribution of synaptic input areas. Correspondingly, we found it useful to inspect the connectome using three compact matrices, based on different normalization schemes: 1) A global normalization that shows the fraction of synaptic area dedicated to a particular pre-post cell type pair, normalized by the total synaptic area of the connectome (Fig. 6a). 2) Output normalization, showing the fraction of synaptic output to each cell type, normalized by total output area over cell types in each row (Fig. 6b). This shows the preferences of a cell type toward connecting with other types. 3) Input normalization, showing the fraction of synaptic input area normalized by total synaptic input from all cell types in each column (Fig. 6c), showing the preference of a cell type for different input types. In combination, these matrices reveal a plethora of previously undescribed circuit organization within the songbird BG (Fig. 6a-c).

In particular, we have identified several distinct networks providing recurrent and feedforward inhibitory control of the direct pathway. As described above, the direct pathway, and not the indirect pathway, provides the predominant contribution to the BG output (Fig. 6d). A notable observation is the recurrent inhibition in the GPi neuron population, involving direct connectivity between GPi neurons (Fig. 6e). Moreover, a significant GPi to STN connectivity, described here, as well as STN to GPi connections, also previously described^13,21^, provide an additional pathway for net mutual inhibitory interactions between GPi neurons.

Second, a feedforward inhibitory pathway is provided by INT1 neurons at the pallidal level. These neurons receive surprisingly strong cortical input from LMAN, which conveys action signals. INT1 neurons project primarily to GPi neurons and provide a feedforward inhibitory path that may serve to balance the hyperdirect LMAN to STN projection (Fig. 6f).

Third, an additional set of feedforward and feedback pathways are centered around regulation of MSNs (Fig. 6g). The strong projection of GPe neurons onto INT2 and INT3 interneurons provides recurrent inhibitory feedback for MSNs. Additionally, the HVC to INT2/3 projection implements feedforward inhibition that may balance the cortical, predominantly HVC, inputs onto MSNs.

## Discussion

Based on a synaptic resolution EM dataset of zebra finch Area X, a circuit essential for song learning, we performed the first in-depth connectomic analysis of vertebrate BG pathways. Our approach uses automated connectome reconstructions, rather than relying on extensive manual proofreading. Systematic manual inspection of reconstructed neurons revealed a relatively small number of apparent erroneous reconstructions (Extended Data Table 3). This highlights the progress of automated connectome analysis over the past decade^41,42,63^ and stands in contrast to recently released full connectomic datasets of the fly, which required massive proofreading efforts^64,65^. It should be noted that our automated neuron reconstructions are not perfect, and different structures suffer from different error rates, e.g., thin axons are more frequently fragmented (Extended Data Fig. 4) than large caliber dendrites. This also means that the presented connectivity data depends on filtering criteria (Extended Data Fig. 2), particularly for putative HVC and LMAN axons. Similarly, cell-type identification is more reliable for neurons with a soma within Area X, in contrast to incoming axon fragments, which are harder to classify. For example, it has been reported that there is a small thalamic projection to Area X in bengalese finches^66^, as observed for the mammalian striatum^67^ but these did not appear as a separate cluster in our analysis. These remaining uncertainties will ultimately be resolved by much larger connectomes, ideally spanning the entire songbird brain, which will make it trivial to resolve and quantify neuronal identity unambiguously.

Across different vertebrate species, connectivity through the basal ganglia has been described in terms of three primary pathways: the direct, indirect and hyperdirect pathway ^3,68^. In songbird, previous work in Area X described separate GPi and GPe neuron populations, supporting the existence of an indirect, as well as a direct pathway^38^. Other early work suggested the existence of local excitatory neurons^39^, further supporting the existence of an indirect, and possibly hyperdirect pathway. Our findings allow us to not only trace these pathways synaptically, but make quantitative statements about their relative strengths. As MSNs are part of both the direct and indirect pathways, we begin such quantification by defining which MSN neurons belong to each pathway.

In the mammal, MSNs specific to the direct or indirect pathways show differences in morphology, D1 or D2 dopamine receptor expression, and output connectivity^14,51^. Indirect pathway MSNs (those expressing only D2 receptors) and MSNs that co-express D1 and D2 receptors^69^, exclusively target the GPe, while direct pathway MSNs (those expressing only D1 receptors) project to the GPi with collateral projections to the GPe^26^. In our data, most individual MSNs (> 70%) project to both the GPe and GPi, while only a tiny fraction are GPe-specific (3%). Altogether, at least 94% of MSNs project to GPi neurons, consistent with a highly predominant direct pathway. Additionally, MSN connectivity to GPe and GPi was not associated with differences in MSN morphology. Thus MSNs in our dataset are not separable into distinct subpopulations either by connectivity or morphology. Previous work on D1 and D2 receptor expression in Area X MSNs did find differential expression of D1 and D2 receptors, although with a large fraction of neurons expressing both D1 and D2 receptors^40,70^. How these findings relate to the observed connectivity in our dataset remains to be elucidated.

It has remained an important open question whether there is an analog of the indirect and hyperdirect pathways, and their associated STN neurons, in Area X^39^. Electrophysiological evidence^39^, as well as evidence from gene expression data^40^ showed that excitatory neurons likely exist inside of Area X, and make glutamatergic synaptic connections onto GPi neurons. Another possibility is that song-related STN neurons exist outside Area X. Indeed, singing-related activity in a subthalamic nucleus outside of Area X has been observed in zebra finches ^71^. Our results confirm a population of neurons in Area X that have a morphology resembling mammalian STNs^53–55^. These make putative excitatory synapses onto GPi cells, supporting the existence of indirect and/or hyperdirect pathways entirely within Area X. However, there may be some significant differences between this pathway within Area X and that observed in mammals. For example, GPe to STN and GPe to GPi projections, key links of the indirect pathway, are relatively weak, suggesting that the hyperdirect (rather than the indirect) pathway plays a crucial role in counterbalancing the direct pathway.

Given the large number of neuron reconstructions, our data allowed us to perform an extensive analysis of BG neuronal morphology and connectivity (Fig. 1, 6). We were able to reconstruct, at ultrastructural resolution, hundreds of neurons belonging to rare interneuron types (Fig. 5, Extended Data Fig. 12, 13). For example, in our data, three types broadly resemble the FS interneuron from the mammalian striatum^34^. While sparse electrophysiological recordings and labeling studies found only one FS-type in Area X^34,36^, single nucleus RNA sequencing data suggested a large number of interneuron clusters^40^, foreshadowing new neuron types found our own connectomic results. In our data, the three types that are clearly distinct morphologically (Fig. 5a-c), also differ substantially in their incoming and outgoing connectivity profiles (Fig. 5d, 6). Potential mammalian analogues for INT2 and INT3 include FS^72^, neurogliaform interneurons^73^ or arkypallidal neurons^74^, all of which are GABAergic neuron types that strongly innervate MSNs. Based on the observation of autapses and the feedforward inhibition of MSNs from HVC via INT2/3, the INT3 may be most similar to mammalian FS interneurons^75^. Given their preferential targeting of pallidal neurons, INT1 neurons likely correspond to pallidal local interneurons. Future connectomic, physiological and molecular expression studies will be needed to clarify how similar the INT1,2 and 3 neurons are to their mammalian analogues.

Another novel finding in our data is the strong lateral inhibition between neighboring GPi neurons. This mutual inhibition acts through direct somatic contacts between GPi neurons, and through the strong mutual connectivity between STN and GPi (Fig. 6e). Our data thus raise the question of what functional or computational role these connections may play: for example, as a competitive interaction between alternative behaviors^76^. A notable feature of this recurrent inhibition in Area X is its possible regulation by the feedforward inhibition from LMAN, mediated by the population of INT1 interneurons. More specifically, LMAN provides direct inputs onto STN neurons (‘hyperdirect pathway’) while INT1 neurons receive a collateral of this LMAN input, and in turn inhibit local GPi neurons.

In contrast to the classical view of GPe neurons as a feed-forward ‘no-go’ pathway, in Area X these neurons appear to serve a role in recurrent inhibition of MSNs through the population of INT2 and INT3 interneurons (Fig. 6g). More specifically, activation of GPe-projecting MSNs would inhibit GPe neurons, releasing INT2,3 interneurons to fire, thus providing broad inhibition of the local MSN population. Such inhibition could serve to sharpen MSN responses in both space and time, or produce winner-take-all dynamics. If activation of GPe-projecting MSNs is controlled by learning at HVC inputs, this would provide a temporally specific tunable input that controls such lateral sharpening. These examples of recurrent connectivity loops, alongside the canonical BG circuitry in the song system, likely play important roles in learning.

Further investigation into the function of these feedback loops may provide a new perspective on mammalian BG neuroscience. With the advent of modern automated methods for the acquisition and analysis of connectomes^41,42^, we expect a large number of vertebrate BG connectomes in the near future, which will allow in-depth comparisons across - and within species. These might be particularly informative in the context of how neurological disorders rewire the circuitry, and we hope that the first synaptic resolution vertebrate BG connectome can serve as a powerful comparison point.

## Material and Methods

### Animal Handling

The dataset was collected from an adult, male zebra finch (> 120 dph). Perfusion and animal handling were performed by members of the Fee lab at MIT in accordance with NIH guidelines, and reviewed and approved by the Massachusetts Institute of Technology Committee on Animal Care.

### Sample preparation

Perfusion was performed by members of the Fee Lab, MIT with 0.07 M cacodylate buffer (CB), 0.14 M sucrose, 1% GA, 1% PFA, as described earlier. In short, to open the blood-brain barrier, high pressure was used to apply the fixative solution^77^. After extraction, the brain was sectioned sagittally at 300 µm using a vibratome (Leica VT1000S). Area X was identified using a stereomicroscope and a slice with a cross section of Area X, was taken and postfixed in the fixative solution overnight. After several washes with 0.15 M CB, osmium staining was performed according to the ROTO protocol^78^. In brief, the sample was stained with a reduced osmium solution (2% OsO4, 2.5% potassium hexacyanoferrate (II), 0.15 M CB) for 2 hours, at room temperature (RT). After several rinses with 0.15 M CB, the sample was immersed in 1% thiocarbohydrazide (TCH) at 58 °C for 1 hour. After multiple rinses in ddH2O, also between the following steps, the sample was first immersed in 2% OsO4, RT, 2 hours, 2% uranyl acetate, 53 °C, 2 hours, then 0.02 M lead aspartate, 53 °C, 2 hours. The sample was then dehydrated through a graded series of cooled ethanol concentrations (70%, 100%, 100%, 100% for 10, 15, 10, 10 min). For embedding the sample was immersed two times in 100% propylene oxide, then with a 50/50 propylene oxide/ epoxy mixture and then with 100% epoxy. The epoxy was 812-replacement (hard mixture, Serva). After infiltration, samples were cured at 60°C for 48h. The samples were trimmed, gold-coated to improve conductivity and then smoothed with an ultramicrotome (Leica Ultracut microtome).

### Electron Microscopy Imaging

The sample was acquired with serial block-face electron microscopy, with a microscope equipped with a Zeiss UltraPlus field emission cathode and a custom-built in-chamber microtome^44^. Images were acquired in high vacuum with a backscatter detector, 2 kV electron landing energy and a scan rate of 5 MHz, an electron current of 1 nA, a lateral resolution of 10 x 10 nm and a cutting thickness of 25 nm. Images were first registered coarse^79^, and then fine aligned, including elastic alignment, by A. Pope. The size of the volume containing the dataset is 256 x 256 x 384 µm^3^.

### Data Processing

Data were processed as described in SyConn2^42^ with some additional adjustments. The neurite reconstructions and compartment predictions are the same but mitochondria, vesicle clouds and synapses were processed again using the SyConn2 pipeline but extended ground truth. The ground truth consisted of 51 cubes sized 300 x 300 x 150 voxels each, where mitochondria and synaptic junctions were annotated. For vesicle clouds the ground truth was not updated. For mitochondria additionally six cubes with the same size that included myelin were annotated. To make sure the individual mitochondria would not be predicted together, a binary erosion (scipy.ndimage.binary_erosion) was performed for two iterations and a border_value of 1. Annotations were performed by trained student annotators and refined by AR and JK. The training and prediction of the new organelle segmentations was done by MJ.

For synapses, a random forest classifier (RFC), was used to predict synapse probability^42,43^. The RFC was trained on 330 synapses with 30 randomly selected synapses from each cell type (INT1-3 were still divided into FS and NGF only), for neuronal cell types residing in Area X, 15 synapses were incoming and 15 were outgoing. Only axo-dendritic and axo-somatic synapses were selected. They were manually labeled true/false by four expert annotators including AR. The annotators were given only information about the synaptic coordinates to verify with SyConnWeb where information about cell type but not mesh area was available. The final label was assigned by majority vote, in case of a tie AR’s vote resulted in the final label. In the end, this resulted in 234 true and 96 false synapses. After cross-validation the resulting F1-score was 0.88, the non-synaptic score was 0.68, and the accuracy was 0.82 (see Extended Data Table 1). For cell typing a supervised machine learning approach was used that runs on point clouds of the mesh of the cells, as described previously^42^.

Cell types were based on morphological descriptions in the literature, similar to the ones described before^42^, but with extended cell types (see Table 1). Additional cell types added were astrocytes, oligodendrocytes, microglia and migratory neurons, while previous FS and NGF class neurons were split into INT1-3. Previous FS ground truth cells were relabeled INT1, while previous NGF ground truth cells were sorted into INT2 and INT3 categories based on manually set thresholds. These thresholds are based on clusters in axonal mitochondria volume density, soma diameter, axon median radius, and spine density. Soma diameter was estimated from the compartment prediction and the median distance to the median soma vertex coordinate was used. Thresholds were set at 0.025 µm^3^/µm for axonal mitochondria volume density, 0.11 µm for axon median radius, 11.5 µm for soma radius, 0.025 1/µm for spine density (see Fig. 5, Extended Data Fig. 11). Cells that did not fit into one category for all thresholds were removed from the ground truth and additional cells were added. Further information on the calculation of these parameters is provided in the following sections.

The ground truth for GPe and GPi cells was based on previous descriptions for Area X pallidal-like cells (Table 1)^34,50^. Cells with smaller somata and more locally branching axons were used as ground truth for GPe, while cells with larger somata and more extensively branching axons that leave the dataset (as expected for cells projecting outside of Area X) were used as ground truth for GPi. To demonstrate that this initial manual classification resulted in morphologically distinct classes, a four-dimensional PCA was performed using the following parameters: axonal mitochondria density, axon median radius, axon myelin fraction, and soma diameter (see Methods: Morphological Analysis). The ground truth for projecting axon types (HVC, LMAN, DA) was extended, and to avoid potential bias related to fragment length, it was ensured that different fragment lengths of each class were included (see Extended Data Fig. 4b). A total of 393 cell types were used as ground truth, the number per cell type is summarized in Extended Data Table 4. The model used for prediction was trained on the entire ground truth without myelin information and with 50k points, 20 µm context and a redundancy of 20, meaning 20 predictions were made per cell (small fragments sampled with replacement). A 10-fold cross-validation of the model was performed separately on three different splits of the data into training and validation sets, as previously described^42^. The results of the 10-fold cross-validation are reported in Extended Data Table 5 and 6 with results for three validation splits.

The dataset with labeled cell IDs, mitochondria, vesicle clouds and synapses is available at syconn.esc.mpcdf.mpg.de by selecting ‘j0251’ as dataset and ‘j0251_72_seg20210127_syn_20220811_celltypes_20230822’ as version. Cell IDs used in figures are visible with the provided links and can be further inspected by pasting the ids into the field under ‘Seg.’. Coordinates specified in figure captions can be pasted into the coordinate field in the upper left to further visualize the raw data.

### Data quality assessment

#### Synapse quality

To assess the synapse quality, the distribution of synapse probabilities assigned by the RFC was plotted (see Extended Data Fig. 1). In addition, an expert annotator, who also participated in the ground truth verification, manually checked 330 synapses from different synaptic probabilities (categories: 0 - 0.2, 0.2 - 0.4, 0.4 - 0.6, 0.6 - 0.8, 0.8 - 1). 180 of these synapses were filtered by only one synaptic partner being an axon (axo-axonic synapses were allowed). For each neuronal cell type, three synapses were randomly selected in each probability category. The other 152 synapses came from synapses filtered with a minimum synaptic area of 0.1 µm, only axo-dendritic and axo-somatic synapses were allowed and also only synapses between neuronal cell types with a minimum pathlength (50 µm for projecting axons, 200 µm for other neuronal cell types). These were the same criteria used for the connectivity analyses. There, also three synapses were randomly selected from each presynaptic cell type and each category. In the lowest probability bin (0 - 0.2), there were only eight synapses in total, all of which were included. To avoid potential bias, the annotator was given only the synaptic coordinates and the two cell IDs of pre and postsynaptic partners that could be evaluated in SyConnWeb (where information about cell type is available), but no information about synaptic probability or synaptic area. After initial evaluation by one annotator, the synapses classified as ‘unclear’ were reviewed by AR for final classification.

#### Neuron reconstruction

The quality of the neuron reconstructions was assessed in several ways. First, the estimated run length of the automated segmentations was measured^41^, and second, manual proofreading on neurons was performed to estimate error rates for different cell types.

Two different sets were used for the run length measurements. One consisted of 50 selected cells with a soma in the volume that were manually skeletonized and improved over 27 iterations. The second one was done in a densely reconstructed and proofread 25 µm cube (see Extended Data Table 3 for results).

To investigate the number of mergers, 500 randomly selected MSNs with an axon and dendritic pathlength of 200 µm each were manually reviewed by an expert annotator. 6 cells (1.2%) had a merger with either an axon fragment or another cell (see Extended Data Table 3). Additionally, all cells of the cell types GPe, GPi, LTS, STN, TAN that met the aforementioned criteria (245 in total) were manually reviewed for mergers and the presence of axon, soma and dendrite. For INT1-3, the dense arborizations made manual reviews very difficult, so a random subset of 10 cells each was evaluated for mergers. These were manually reviewed independently by two expert annotators. Cases where the opinions differed (3 cells) were reviewed again.

To analyze the fragmentation of projecting axon classes, the skeleton length of these was analyzed.

Axon fragmentation occurred in all projecting axon classes, with a significant proportion of axons of all classes being shorter than 10 µm (see Extended Data Fig. 4c). Unless stated otherwise, only fragments >= 50 µm were included in the analyses, as detailed below.

To see how fragmentation might affect analyses of nearly complete cells, the proportion of nearly complete cells with an axon or dendrite was calculated based on both the number of cells/fragments and the pathlength in the corresponding compartment (see Extended Data Table 7).

### Compartment identification quality

Compartment predictions were the same as used previously^42^. For additional verification, some examples were reviewed manually (Extended Data Fig. 3).

### Filtering for complete cells

Neuronal cell types located in Area X (MSN, STN, TAN, GPe, GPi, LTS, INT1-3) were filtered to ensure sufficient completeness for analyses. This means that all cells used had to have at least one skeleton node predicted as axon, soma and dendrite (compartment prediction as in ^42^ with a skeleton length of axon and dendrite of 200 µm each). Additionally, all cells of the cell types GPe, GPi, LTS, STN and TAN that met this criterion were manually reviewed. Cells that did not contain all three compartments or cells that were fused with a fragment of at least 50 µm pathlength or contained at least one synapse were excluded (24 in total, see Extended Data Table 3). Due to the high number of MSNs and the dense axonal branching in INT1-3, not all cells of these cell types were manually reviewed but a random subset was selected to estimate the error rate (see ‘Neuron reconstruction’, Extended Data Table 3). For comparison, the connectivity matrices in Fig. 6 were also computed without excluding any cells from GPe, GPi, LTS, STN and TAN (see Extended Data Fig. 2a).

This resulted in 8576 cells used for analyses. In the analyses of morphological differences between MSNs (Fig. 2), an additional analysis was performed to filter out all MSNs that did not have a complete dendrite. Since dendrites are thicker than axons, the risk of being cut off in the middle of the dataset due to segmentation errors is very low. To identify cut-off dendrites, the skeleton of a dendrite was checked to see if it was within 7 µm of the dataset boundaries. If so, the MSN cell was excluded from this analysis. The 7 µm threshold was chosen as a threshold from the dataset boundary and not from the segmentation boundary. Due to the shift in alignment, the effective size of the dataset (256 x 256 x 384 µm³) is not perfectly cubed and therefore different than the bounding box of the dataset displayed on SyConnWeb (syconn.esc.mpcdf.mpg.de) or used in the data processing (271 x 271 x 387 µm³). To bridge the distance between the bounding box and the start of the segmentation, 7 µm were selected (e.g. visible at this coordinate: 27125, 10327, 5365). In rare cases, the distance between the segmentation and the dataset boundary is larger (e.g. at this coordinate: 23692, 25488, 7765).

Cell types belonging to projecting axons were generally filtered with 50 µm total skeleton length. This is because axons are thinner and therefore more fragmented than dendrites. The average axonal fragment with at least one synapse is about 10 µm in length (see Extended Data Fig. 4). Applying a filter criterion of at least 200 µm, similar to that used for full cells, would have excluded the majority of axonal fragments and thus risked underestimating their influence of the connectivity matrix. Including all fragments, on the other hand, would overestimate their influence on connectivity compared to the full cells and include potential misclassified axon types, as cell typing on short fragments is more uncertain than on longer ones. The 50 µm were therefore chosen as a compromise. The connectivity matrix was also calculated for a threshold of 0 µm and 200 µm for projecting axons, and once with 50 µm for full cells for full cells and is available in Extended Data Fig. 2b-d.

### Morphological analyses

Analysis of axonal or dendritic length refers to the skeleton length for each compartment, based on the compartment prediction mapped to the cells^42^.

Soma radius was calculated by using the median distance to the center of the soma. The center was the median coordinate from all vertices of the cell predicted as soma. The median axon radius was calculated from the diameter of the skeleton nodes that were predicted to be axons by the compartment classifier.

Axonal mitochondria volume density was calculated by the sum of mitochondrial voxel size times the product of the voxel size (10 nm x 10 nm x 25 nm) relative to the skeleton length of the axonal compartment. The mitochondria were mapped to the compartment by using scipy.spatial.cKDTree to map the mitochondria to the nearest skeleton node and selecting only those that were closest to an axonal value. The mitochondria were mapped only by their representative coordinate, which could result in large mitochondria that straddle the soma-axon boundary being mislabeled as completely in only one of the two compartments.

PCA (see Fig. 2e, 5c) was performed using the PCA library from sklearn.decomposition. Since the PCA used values of different scales, such as mitochondrial density and soma diameter, the values were standardized to z-scores using sklearn.preprocessing.StandartScaler. The PCA was then applied using one principal component. For the PCA in Fig. 2e for GPe and GPi, the parameters used were: axonal mitochondrial volume density, axon median radius, axon myelin fraction and soma diameter. For the PCA in fig. 5c, parameters were: soma diameter, axonal mitochondria density, axon median radius and dendritic spine density.

For the UMAP (Extended Data Fig 6), the umap-learn library was used. Similar to the PCA, the data was standardized with sklearn.preprcessing.StandartScaler. The UMAP was performed on an 18 dimensional feature vector consisting of the following parameters: axon length, axon surface area, axon median radius, dendrite length, dendrite surface area, dendrite median radius, soma diameter, soma surface area, cell volume, axonal, dendritic and soma mitochondrial volume density, vesicle cloud volume density and the synaptic density as sum of synaptic area divided by axon, dendrite, and soma surface area. This feature vector was found to separate most neuron classes well (see Supplementary Information and Extended Data Fig. 6 b,c). While the mitochondria density for axon and dendrite were calculated as the sum of mitochondrial volume per skeleton pathlength, the soma mitochondrial volume density was calculated as the sum of mitochondrial volume density relative to the estimated soma volume density. The soma volume density was calculated with the soma diameter, estimated as described above (see Data Processing).

### Analyses of connectivity

The analysis of synaptic connectivity was performed only on synapses with a synapse probability of 0.6 as assigned by an RFC (see ‘Data Processing’) and a minimum synapse size of 0.1 µm². Synapse size was calculated as the synaptic mesh area divided by 2.

Synapses were only included in the analysis if they were axo-dendritic or axo-somatic. For this, the compartment prediction which was mapped to the mesh vertices of the cell and averaged over 10000 vertices (key = ‘axoness_avg10000’)^42^ was mapped to the synapses using a scipy.cKDTree. Synapses that were not between an axon and a dendrite or soma were then excluded.

For compartment specific connectivity, the ‘axoness_avg10000’ key was used to determine whether the synapse was connected to a dendrite or a soma. To distinguish between dendritic shaft, spine neck and spine head the prediction of a CNN learning dendritic morphology was used and mapped from the mesh vertices to the nearest synapse (key = ‘spiness’)^42^. The values shown in Fig. 6 represent the median percentage of the sum of synapse size on each compartment shown and therefore do not add up to 100%.

For the global connectivity matrix, synapse sizes from all cells of a presynaptic cell type to all cells of a postsynaptic cell type were summed up and divided by the sum of all the synapses in the dataset that were used for analysis. For the incoming connectivity matrix, values were normalized based on the sum of connectivity from all synapses to that cell type (columns sum to 100%). For the outgoing connectivity matrix, values were normalized based on the sum of connectivity from all synapses to this cell type (rows add up to 100%).

Autapses were not registered as synapses in the SyConn framework. Therefore, the identification of autapses was done manually. To determine the fraction of each cell type with at least one autapse within 10 µm of the soma, all cells from the GPe, GPi, STN, LTS, TAN, INT1, INT2, and INT3 datasets with complete somata were examined. Additionally, 50 randomly selected MSNs were also reviewed.

### Simulation of MSN-GPe/GPi connectivity

To test whether there was a preference for MSN to connect to GPe or GPi, the observed ratio of GPe to GPi was compared to shuffled data. The GP ratio was calculated as GPi/(GPe + GPi) area or number of synapses.

To shuffle the data, the synapses and their targets remained the same but their association with the MSN presynaptic cell ids changed. In this way, the number and synaptic area to each target remained the same as in the observed data, where 75% of the synaptic area of MSN-GP synapses goes to GPi and 73% of the synapses go to GPi.

This shuffling was performed for 100 iterations. After each iteration, the number of cells per bin was calculated along with the mean and 95% CI of the shuffled data. The number of bins for both shuffled and observed data was 30.

To calculate the number of GPi-specific MSN cells, all cells with a GP ratio of at least 0.9 were counted for the observed data and each shuffle iteration.

### Cell-specific recurrent inhibition analyses

If there is a cell type that receives both input from and output to the same cell type, there is a possibility that the interaction is between individual cells. To test this, first the summed synaptic area from each cell was calculated to and from each cell of the second cell type. To see how much overlap there was between incoming and outgoing cells, two measurements were made: first, the overlap in synaptic area was calculated, as a ratio for both the incoming and the outgoing sides. This was done by dividing the sum of the synaptic area of cells that were on both the incoming and outgoing sides by the total sum of incoming synapses to that cell by that specific cell type (accordingly also for outgoing, see Extended Data Fig. 10b-g). The second one was a binary score of whether the contact with the largest incoming summed synaptic area was also the contact to which the largest output went. This was called: strongest partner reciprocity (see Extended Data Fig. 10h). To get an estimate for the whole population, the median of the incoming and outgoing ratio for both cell types that were part of the connection was calculated together with the fraction of cells that had a strongest partner reciprocity. This analysis was performed for STN-GPi, STN-GPe and MSN-TAN connections. Cells that did not both receive input and project to the other cell type were excluded (cell numbers in Extended Data Table 10).

### Data visualization

Plots were visualized using the seaborn and matplotlib libraries. For cell renderings, cell meshes that were created with the open_3d.TriangleMesh library were exported using syconn.proc.meshes.write_mesh2kzip which uses the PlyData library to export them as .ply files^42^. These files were then imported and rendered in Blender.

### Statistical Analysis

All statistical analyses were performed in Python using the scipy.stats package. Only nonparametric tests were used. These include the calculation of the Spearman rank correlation coefficient to test correlations and the Kruskal-Wallis H-test for analyses with more than two cell types with a Wilcoxon rank sum test as a post hoc test. P values < 0.005 were considered significant.

## Acknowledgements

We thank Winfried Denk for generous intellectual and material support of this project at the Max Planck Institute for Biological Intelligence. We thank Lorenz Hüdepohl and Christian Guggenberger for their support at the MPCDF in Garching. We are grateful to Delta Schick for the manual verification and identification of glial cell types, and to Ben Scott and Naomi Shvedov for the identification and verification of migratory neurons. We also thank Julia Kuhl for her assistance with figure preparation. Our thanks go to Hashir Ahmad, Philipp Schubert, and Andrei Mancu for help with SyConn2 processing. We thank Ben Scott, Jesse Goldberg and Maya Ketzev for feedback on the manuscript. We further thank Riccardo Morbio, Delta Schick, and Laura Werner for the manual verification of synapses used in the RFC ground truth. We especially thank Riccardo Morbio for additionally evaluating RFC predictions on synapses and reviewing cells for mergers and autapses. We are also grateful to the annotators who contributed to the synapse and mitochondria ground truth: Julian Hendricks, Delta Schick, Riccardo Morbio, Katyayni Ganesan, Deniz Üreyener, Laura Werner, Merve Cetiner, Gizem Karabiyik, Yona Perstat, Ata Kan, Dominik Melzer, Akshaya Rajan, and Maria Nikitina. Finally, we thank Delta Schick and Julian Hendricks for reviewing the annotations. This work was funded by the Max Planck Society, Google Research, grant NIH RF1 MH117809 and UKRI.

## Code availability

Data was prepared using the SyConn2 pipeline^42^, [SyConn repository](https://github.com/StructuralNeurobiologyLab/SyConn.git)^42^ and analysed with code available on [github](https://github.com/StructuralNeurobiologyLab/areaX-connectomic-analysis.git).

## Data availability

The dataset is available on SyConn Web (syconn.esc.mpcdf.mpg.de) under the dataset ‘j0251’ and the version ‘j0251_72_seg_20210127_agglo2_syn_20220811_celtypes_20230822’.

## Ethics declarations

Competing interests: JK holds shares of ariadne.ai ag.

## Author contributions

JK collected the volume EM dataset. AR and JK annotated training data and coordinated data annotation with contributions from MJ. JK and MJ processed and aligned the volume EM data. AR and JK coordinated and performed the connectome extraction with SyConn v2, based on EM segmentations by MJ. AR analysed the connectome and prepared figures with contributions from JK. AR, JK and MSF wrote the manuscript. JK and MSF conceptualized the study with contributions from AR, JK, MSF and VJ supervised the study and all authors reviewed the manuscript.

## References

1. Hollerman, J. R., Tremblay, L. & Schultz, W. Involvement of basal ganglia and orbitofrontal cortex in goal-directed behavior. Prog. Brain Res. 126, (2000).

2. Shipp, S. The functional logic of corticostriatal connections. Brain structure & function 222, (2017).

3. Grillner, S. & Robertson, B. The Basal Ganglia Over 500 Million Years. Curr Biol 26, R1088–R1100 (2016).

4. Alexander, G. E., DeLong, M. R. & Strick, P. L. Parallel organization of functionally segregated circuits linking basal ganglia and cortex. Annu. Rev. Neurosci. 9, 357–381 (1986).

5. DeLong, M. R. Activity of pallidal neurons during movement. J Neurophysiol 34, 414–427 (1971).

6. Graybiel, A. M., Aosaki, T., Flaherty, A. W. & Kimura, M. The basal ganglia and adaptive motor control. Science 265, 1826–1831 (1994).

7. Pierce, J. E. & Péron, J. The basal ganglia and the cerebellum in human emotion. Soc. Cogn. Affect. Neurosci. 15, (2020).

8. Obeso, J. A. et al. Pathophysiology of the basal ganglia in Parkinson’s disease. Trends Neurosci. 23, S8–S19 (2000).

9. Yager, L. M., Garcia, A. F., Wunsch, A. M. & Ferguson, S. M. The ins and outs of the striatum: role in drug addiction. Neuroscience 301, 529–541 (2015).

10. Lanciego, J. L., Luquin, N. & Obeso, J. A. Functional Neuroanatomy of the Basal Ganglia. Cold Spring Harb. Perspect. Med. 2, (2012).

11. Hardman, C. D. et al. Comparison of the basal ganglia in rats, marmosets, macaques, baboons, and humans: volume and neuronal number for the output, internal relay, and striatal modulating nuclei. J. Comp. Neurol. 445, 238–255 (2002).

12. Albin, R. L., Young, A. B. & Penney, J. B. The functional anatomy of basal ganglia disorders. Trends Neurosci. 12, 366–375 (1989).

13. Nambu, A., Tokuno, H. & Takada, M. Functional significance of the cortico-subthalamo-pallidal ‘hyperdirect’ pathway. Neurosci. Res. 43, 111–117 (2002).

14. Gerfen, C. R. et al. D1 and D2 dopamine receptor-regulated gene expression of striatonigral and striatopallidal neurons. Science 250, 1429–1432 (1990).

15. Le Moine, C. & Bloch, B. D1 and D2 dopamine receptor gene expression in the rat striatum: sensitive cRNA probes demonstrate prominent segregation of D1 and D2 mRNAs in distinct neuronal populations of the dorsal and ventral striatum. J. Comp. Neurol. 355, 418–426 (1995).

16. Ericsson, J. et al. Dopamine differentially modulates the excitability of striatal neurons of the direct and indirect pathways in lamprey. J. Neurosci. 33, 8045–8054 (2013).

17. Chevalier, G. & Deniau, J. M. Disinhibition as a basic process in the expression of striatal functions. Trends Neurosci. 13, 277–280 (1990).

18. Kravitz, A. V. et al. Regulation of parkinsonian motor behaviours by optogenetic control of basal ganglia circuitry. Nature 466, 622–626 (2010).

19. Cui, G. et al. Concurrent activation of striatal direct and indirect pathways during action initiation. Nature 494, 238–242 (2013).

20. Cazorla, M. et al. Dopamine D2 receptors regulate the anatomical and functional balance of basal ganglia circuitry. Neuron 81, 153–164 (2014).

21. Nambu, A. et al. Excitatory cortical inputs to pallidal neurons via the subthalamic nucleus in the monkey. J. Neurophysiol. 84, 289–300 (2000).

22. Hannah, R. & Aron, A. R. Towards real-world generalizability of a circuit for action-stopping. Nat Rev Neurosci 22, 538–552 (2021).

23. Fife, K. H. et al. Causal role for the subthalamic nucleus in interrupting behavior. Elife 6, (2017).

24. Kemp, J. M. & Powell, T. P. The termination of fibres from the cerebral cortex and thalamus upon dendritic spines in the caudate nucleus: a study with the Golgi method. Philos Trans R Soc Lond B Biol Sci 262, 429–439 (1971).

25. Loopuijt, L. D. & van der Kooy, D. Organization of the striatum: collateralization of its efferent axons. Brain Res 348, 86–99 (1985).

26. Wu, Y., Richard, S. & Parent, A. The organization of the striatal output system: a single-cell juxtacellular labeling study in the rat. Neurosci. Res. 38, 49–62 (2000).

27. Johansson, Y. & Silberberg, G. The Functional Organization of Cortical and Thalamic Inputs onto Five Types of Striatal Neurons Is Determined by Source and Target Cell Identities. Cell Rep 30, 1178–1194.e3 (2020).

28. Mooney, R. Neural mechanisms for learned birdsong. Learn. Mem. 16, 655–669 (2009).

29. Brainard, M. S. & Doupe, A. J. What songbirds teach us about learning. Nature 417, 351–358 (2002).

30. Nottebohm, F. & Arnold, A. P. Sexual dimorphism in vocal control areas of the songbird brain. Science 194, 211–213 (1976).

31. Fee, M. S. & Goldberg, J. H. A hypothesis for basal ganglia-dependent reinforcement learning in the songbird. Neuroscience 198, 152–170 (2011).

32. Perkel, D. J. Origin of the anterior forebrain pathway. Ann N Y Acad Sci 1016, 736–748 (2004).

33. Jarvis, E. D. Learned birdsong and the neurobiology of human language. Ann. N. Y. Acad. Sci. 1016, 749–777 (2004).

34. Farries, M. A. & Perkel, D. J. A telencephalic nucleus essential for song learning contains neurons with physiological characteristics of both striatum and globus pallidus. J. Neurosci. 22, 3776–3787 (2002).

35. Person, A. L., Gale, S. D., Farries, M. A. & Perkel, D. J. Organization of the songbird basal ganglia, including area X. J. Comp. Neurol. 508, 840–866 (2008).

36. Goldberg, J. H. & Fee, M. S. Singing-related neural activity distinguishes four classes of putative striatal neurons in the songbird basal ganglia. J. Neurophysiol. 103, 2002–2014 (2010).

37. Goldberg, J. H., Adler, A., Bergman, H. & Fee, M. S. Singing-Related Neural Activity Distinguishes Two Putative Pallidal Cell Types in the Songbird Basal Ganglia: Comparison to the Primate Internal and External Pallidal Segments. J. Neurosci. 30, 7088–7098 (2010).

38. Farries, M. A., Ding, L. & Perkel, D. J. Evidence for ‘direct’ and ‘indirect’ pathways through the song system basal ganglia. J. Comp. Neurol. 484, 93–104 (2005).

39. Budzillo, A., Duffy, A., Miller, K. E., Fairhall, A. L. & Perkel, D. J. Dopaminergic modulation of basal ganglia output through coupled excitation-inhibition. Proc. Natl. Acad. Sci. U. S. A. 114, 5713–5718 (2017).

40. Xiao, L. et al. Expression of FoxP2 in the basal ganglia regulates vocal motor sequences in the adult songbird. Nat. Commun. 12, 1–18 (2021).

41. Januszewski, M. et al. High-precision automated reconstruction of neurons with flood-filling networks. Nat. Methods 15, 605–610 (2018).

42. Schubert, P. J. et al. SyConn2: dense synaptic connectivity inference for volume electron microscopy. Nat. Methods 19, 1367–1370 (2022).

43. Dorkenwald, S. et al. Automated synaptic connectivity inference for volume electron microscopy. Nat. Methods 14, 435–442 (2017).

44. Denk, W. & Horstmann, H. Serial block-face scanning electron microscopy to reconstruct three-dimensional tissue nanostructure. PLoS Biol. 2, e329 (2004).

45. Luo, M., Ding, L. & Perkel, D. J. An Avian Basal Ganglia Pathway Essential for Vocal Learning Forms a Closed Topographic Loop. J. Neurosci. 21, 6836–6845 (2001).

46. Gagnon, D. et al. Striatal Neurons Expressing D and D Receptors are Morphologically Distinct and Differently Affected by Dopamine Denervation in Mice. Sci. Rep. 7, 41432 (2017).

47. Chuhma, N., Tanaka, K. F., Hen, R. & Rayport, S. Functional connectome of the striatal medium spiny neuron. J. Neurosci. 31, 1183–1192 (2011).

48. Shvedov, N. R. et al. Songbird connectome reveals tunneling of migratory neurons in the adult striatum. in preparation.

49. Kornfeld, J. et al. An anatomical substrate of credit assignment in reinforcement learning. in preparation.

50. Luo, M. & Perkel, D. J. Long-range GABAergic projection in a circuit essential for vocal learning. J. Comp. Neurol. 403, (1999).

51. Gertler, T. S., Chan, C. S. & Surmeier, D. J. Dichotomous anatomical properties of adult striatal medium spiny neurons. J. Neurosci. 28, 10814–10824 (2008).

52. Fujiyama, F. et al. Exclusive and common targets of neostriatofugal projections of rat striosome neurons: a single neuron-tracing study using a viral vector. Eur. J. Neurosci. 33, 668–677 (2011).

53. Moriizumi, T., Nakamura, Y., Kitao, Y. & Kudo, M. Ultrastructural analyses of afferent terminals in the subthalamic nucleus of the cat with a combined degeneration and horseradish peroxidase tracing method. J. Comp. Neurol. 265, 159–174 (1987).

54. Canteras, N. S., Shammah-Lagnado, S. J., Silva, B. A. & Ricardo, J. A. Afferent connections of the subthalamic nucleus: a combined retrograde and anterograde horseradish peroxidase study in the rat. Brain Res. 513, 43–59 (1990).

55. Smith, Y., Hazrati, L.-N. & Parent, A. Efferent projections of the subthalamic nucleus in the squirrel monkey as studied by the PHA-L anterograde tracing method. J. Comp. Neurol. 294, 306–323 (1990).

56. Naito, A. & Kita, H. The cortico-pallidal projection in the rat: an anterograde tracing study with biotinylated dextran amine. Brain Res 653, 251–257 (1994).

57. Colonnier, M. Synaptic patterns on different cell types in the different laminae of the cat visual cortex. An electron microscope study. Brain Res 9, 268–287 (1968).

58. Gray, E. G. Axo-somatic and axo-dendritic synapses of the cerebral cortex: an electron microscope study. J Anat 93, 420–433 (1959).

59. Kawaguchi, Y. Physiological, morphological, and histochemical characterization of three classes of interneurons in rat neostriatum. J. Neurosci. 13, 4908–4923 (1993).

60. Lapper, S. R., Smith, Y., Sadikot, A. F., Parent, A. & Bolam, J. P. Cortical input to parvalbumin-immunoreactive neurones in the putamen of the squirrel monkey. Brain Res. 580, (1992).

61. Contant, C., Umbriaco, D., Garcia, S., Watkins, K. C. & Descarries, L. Ultrastructural characterization of the acetylcholine innervation in adult rat neostriatum. Neuroscience 71, 937–947 (1996).

62. Tepper, J. M. et al. Heterogeneity and Diversity of Striatal GABAergic Interneurons: Update 2018. Front Neuroanat 12, 91 (2018).

63. Kornfeld, J. & Denk, W. Progress and remaining challenges in high-throughput volume electron microscopy. Curr Opin Neurobiol 50, 261–267 (2018).

64. Dorkenwald, S. et al. Neuronal wiring diagram of an adult brain. Nature 634, 124–138 (2024).

65. Winding, M. et al. The connectome of an insect brain. Science 379, eadd9330 (2023).

66. Nicholson, D. A., Roberts, T. F. & Sober, S. J. Thalamostriatal and cerebellothalamic pathways in a songbird, the Bengalese finch. J. Comp. Neurol. 526, 1550–1570 (2018).

67. Cowan, W. M. & Powell, T. P. A study of thalamo-striate relations in the monkey. Brain 79, 364–390 (1956).

68. Tanimoto, Y. et al. Transgenic tools targeting the basal ganglia reveal both evolutionary conservation and specialization of neural circuits in zebrafish. Cell Rep 43, 113916 (2024).

69. Bonnavion, P. et al. Striatal projection neurons coexpressing dopamine D1 and D2 receptors modulate the motor function of D1- and D2-SPNs. Nat Neurosci 27, 1783–1793 (2024).

70. Kubikova, L., Wada, K. & Jarvis, E. D. Dopamine receptors in a songbird brain. J Comp Neurol 518, 741–769 (2010).

71. Das, A. & Goldberg, J. H. Songbird subthalamic neurons project to dopaminergic midbrain and exhibit singing-related activity. J Neurophysiol 127, 373–383 (2022).

72. Gittis, A. H., Nelson, A. B., Thwin, M. T., Palop, J. J. & Kreitzer, A. C. Distinct roles of GABAergic interneurons in the regulation of striatal output pathways. J. Neurosci. 30, 2223–2234 (2010).

73. English, D. F. et al. GABAergic circuits mediate the reinforcement-related signals of striatal cholinergic interneurons. Nat Neurosci 15, 123–130 (2011).

74. Mallet, N. et al. Dichotomous organization of the external globus pallidus. Neuron 74, 1075–1086 (2012).

75. Wang, X. et al. Functional Autapses Form in Striatal Parvalbumin Interneurons but not Medium Spiny Projection Neurons. Neurosci Bull 39, 576–588 (2023).

76. Bar-Gad, I. & Bergman, H. Stepping out of the box: information processing in the neural networks of the basal ganglia. Curr Opin Neurobiol 11, 689–695 (2001).

77. Cragg, B. Preservation of extracellular space during fixation of the brain for electron microscopy. Tissue Cell 12, 63–72 (1980).

78. Briggman, K. L., Helmstaedter, M. & Denk, W. Wiring specificity in the direction-selectivity circuit of the retina. Nature 471, 183–188 (2011).

79. Scheffer, L. K., Karsh, B. & Vitaladevun, S. Automated Alignment of Imperfect EM Images for Neural Reconstruction. arXiv [q-bio.QM] (2013).

80. Henselmans, J. M. & Wouterlood, F. G. Light and electron microscopic characterization of cholinergic and dopaminergic structures in the striatal complex and the dorsal ventricular ridge of the lizard Gekko gecko. J. Comp. Neurol. 345, 69–83 (1994).

81. Wildenberg, G. et al. Partial connectomes of labeled dopaminergic circuits reveal non-synaptic communication and axonal remodeling after exposure to cocaine. Elife 10, (2021).

82. Fortune, E. S. & Margoliash, D. Parallel pathways and convergence onto HVc and adjacent neostriatum of adult zebra finches (Taeniopygia guttata). The Journal of Comparative Neurology vol. 360 413–441 Preprint at 10.1002/cne.903600305 (1995).

83. Vates, G. E. & Nottebohm, F. Feedback circuitry within a song-learning pathway. Proc. Natl. Acad. Sci. U. S. A. 92, 5139–5143 (1995).

84. Buchanan, J. et al. Oligodendrocyte precursor cells ingest axons in the mouse neocortex. Proc. Natl. Acad. Sci. U. S. A. 119, e2202580119 (2022).

85. Scott, B. B., Gardner, T., Ji, N., Fee, M. S. & Lois, C. Wandering neuronal migration in the postnatal vertebrate forebrain. J. Neurosci. 32, 1436–1446 (2012).

