## Supplementary Information for "The songbird basal ganglia connectome"

### Data Quality

#### *Synapse quality*

To assess the quality of the synapse prediction and provide an additional filter criterion, a random forest classifier (RFC) is used in the SyConn pipeline[1,2]. The RFC was trained on 330 randomly selected synapses (see Methods for details) to predict a probability between 0 and 1 for each synapse. The synapse probability indicates the fraction of correctly predicted synapses with the corresponding value e.g. a synapse probability of 0.8 indicates 80% of the synapses with this value are indeed true synapses. To test the prediction of the synapse probability, 180 randomly selected synapses were manually reviewed. Using the filter criteria of the data analyses (minimum synapse size = 0.1 µm$^2$, only axo-dendritic or axo-somatic synapses, certain minimum pathlengths, see methods), synapses with low probability disappear (Extended Data Fig. 1 a, b).

For data analysis the synapse probability was used as additional filter criterion. Taken together, synapses with a synapse probability of at least 0.6 had a combined accuracy over 90% (90,27% for all synapses, 93,01% for > 0.1 µm$^2$; bars 0.8, 1 in Extended Data Fig. 10 c, d). Only synapses with a probability of at least 0.6 were included in the analysis.

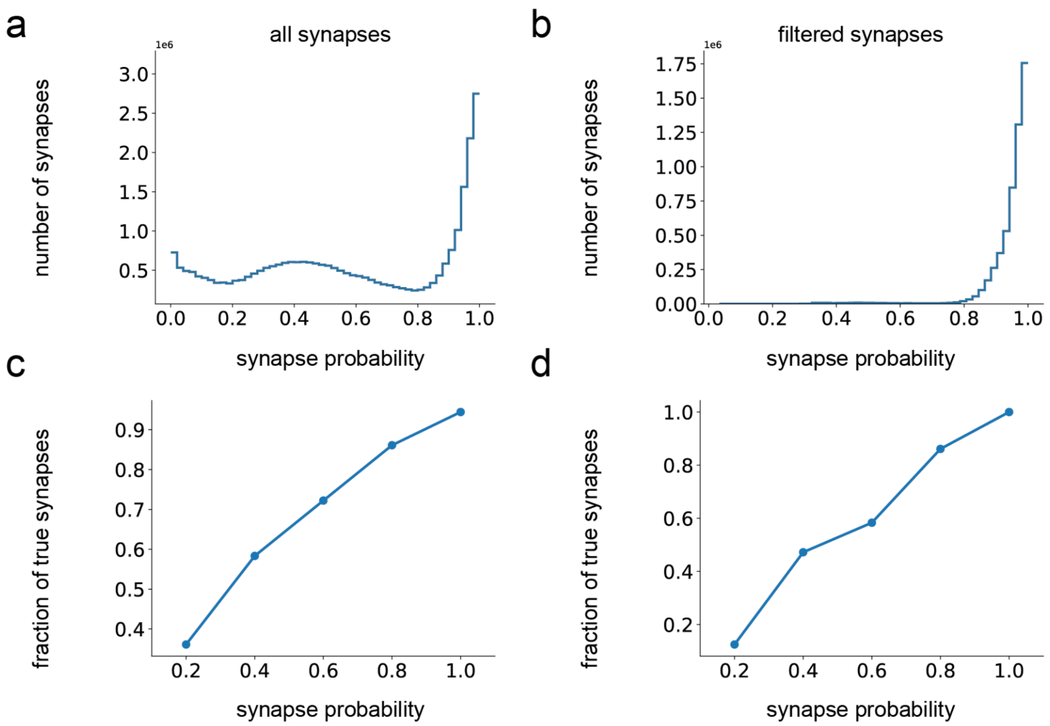


Extended Data Figure 1: Synapse probabilities and RFC evaluation. a RFC synapse probabilities for all synapses in the dataset. b RFC probabilities for filtered synapses with cells from the analysis, axo-dendritic and axo-somatic synapses only, minimum synaptic area = 0.1. c Manual verification of different RFC probabilities. Fraction of true synapses in an evaluation dataset containing 36 synapses of each category (three of each presynaptic cell type). Labels indicate the upper limit of each category, e.g. 0.4 contains synapses with probability 0.2 - 0.4. Synapses were filtered only for the presence of an axon. d Fraction of true synapses in an evaluation dataset similar to c, but with filtered synapses only (see b). Only eight cells with a synapse probability < 0.2 existed in the filtered synapses, so all eight were part of the evaluation dataset. Other categories consisted of 36 synapses (three from each presynaptic cell type).

| | **precision** | **recall** | **F1-score** | **support** |
|---|---|---|---|---|
| **non-synaptic** | 0.70 | 0.67 | 0.68 | 96 |
| **synaptic** | 0.87 | 0.88 | 0.88 | 234 |
| **accuracy** | | | 0.82 | 330 |
| **macro avg** | 0.78 | 0.78 | 0.78 | 330 |
| **weighted avg** | 0.82 | 0.82 | 0.82 | 330 |

Extended Data Table 1: Results of RFC 10-fold cross-validation for synapse classification.

*Neuron reconstruction accuracy evaluation*

We measured the run length in two different sets. One with 50 selected complete cells which was manually skeletonized and improved over 27 iterations and one in a 25 µm cube which was densely reconstructed and proofread (see Extended Data Table 2).

| | **complete cells from whole dataset** | **25 µm cube, all** | **25 µm cube, axons** | **25 µm cube, dendrites** |
|---|---|---|---|---|
| **absolute run length [µm]** | 6906.20 | 31.74 | 22.04 | 63.13 |
| **normalized estimated run length** | 0.693 | 0.699 | 0.650 | 0.765 |

Extended Data Table 2: Segmentation absolute run length and normalized estimated run length for complete cells in the entire dataset and a densely reconstructed and proofread 25 µm cube.

| **cell type** | **number total** | **number erroneous cells** | **number incomplete cells** | **number cells with mergers** | **mergers per mm pathlength** | **number mergers with neurons** | **number mergers with glia** |
|---|---|---|---|---|---|---|---|
| **MSN** | 500 | 6 | 0 | 6 | 0.004 | 6 | 0 |
| **GPe** | 28 | 1 | 1 | 0 | 0.000 | 0 | 0 |
| **GPi** | 58 | 11 | 2 | 9 | 0.020 | 11 | 1 |
| **LTS** | 38 | 1 | 0 | 1 | 0.008 | 1 | 0 |
| **STN** | 108 | 6 | 2 | 4 | 0.006 | 5 | 0 |
| **TAN** | 13 | 5 | 1 | 4 | 0.049 | 5 | 0 |
| **INT1** | 10 | 3 | 0 | 3 | 0.030 | 2 | 1 |
| **INT2** | 10 | 1 | 0 | 1 | 0.018 | 1 | 0 |
| **INT3** | 10 | 1 | 0 | 1 | 0.014 | 1 | 1 |

Extended Data Table 3: Mergers of manually inspected cells by expert annotators. All inspected cells had a soma, axon and dendrite predicted, axon and dendrite with a minimum pathlength of 200 µm each. For MSN, a random subset of 500 cells (out of 8093) was selected, for INT1-3 a random subset of 10 cells, as these cells have extensive axonal arborization, making manual checking time-consuming. For the other cell types (GPe, GPi, LTS, STN, TAN), the entire set of cells was manually inspected. If cells either did not have an axon, soma and dendrite after visual inspection ('incomplete') or were merged with a part of another cells (> 50 µm pathlength or > 1 synapse), the cells were marked as 'erroneous'. Erroneous cells were excluded from the analysis for the cell types, where all cells could be manually inspected (see Methods). To obtain the number of mergers per mm, all mergers with different fragments were counted (some cells had more than one merger) and divided by the skeleton pathlength of all cells of that cell type.

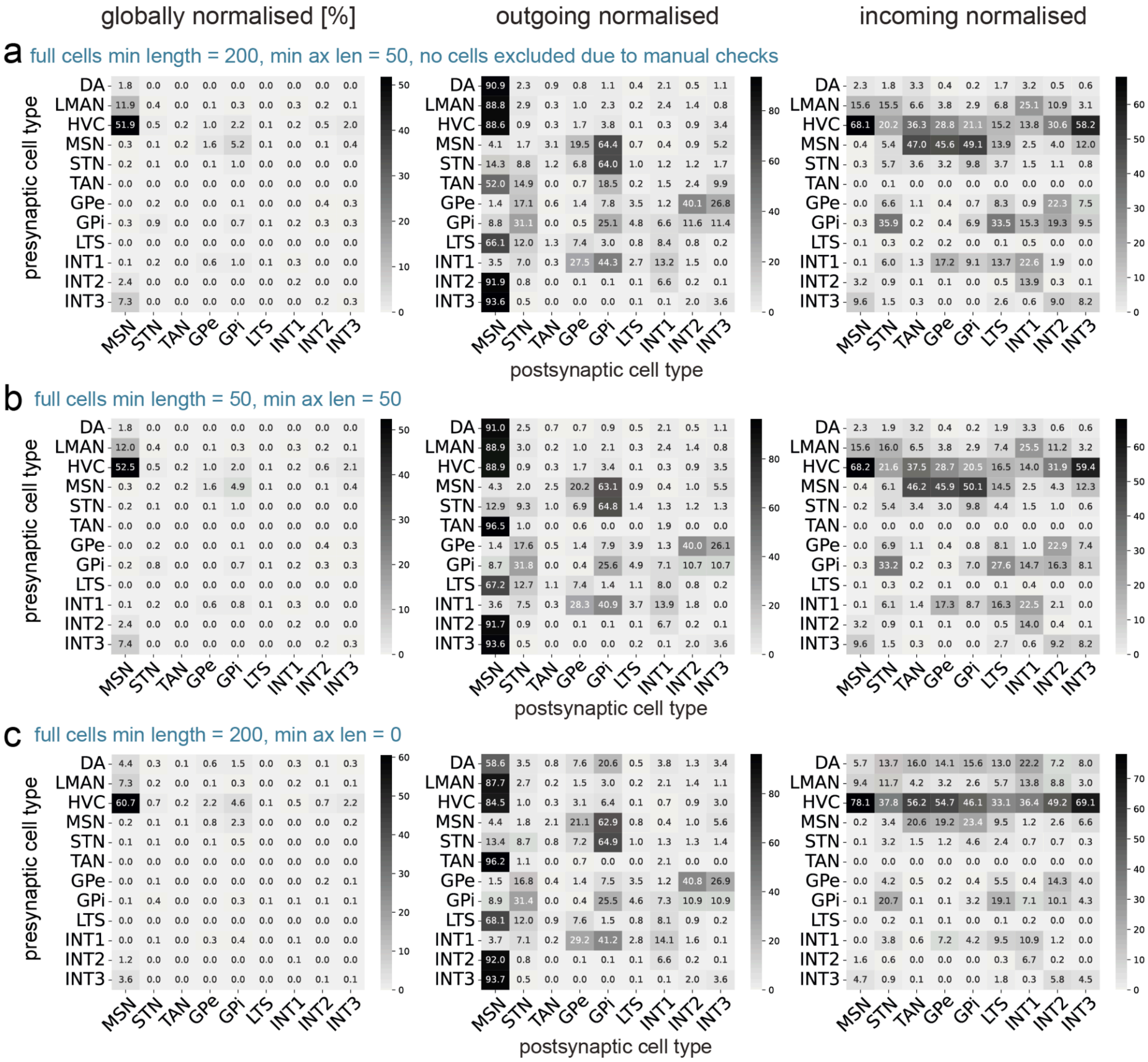


Extended Data Figure 2. Connectivity matrices showing the percentage of the synaptic area with different normalizations as shown in Fig. 6e-g with different filtering criteria. a Matrices with full cells filtered to a minimum axon and dendrite skeleton length of 200 µm each. Projecting axon cell types (DA, HVC,

LMAN) filtered to a minimum skeleton length of 50 µm. Cells that were manually identified as incomplete or mergers of cell types STN, GPe, GPi, TAN, LTS were not excluded (see Extended Data Table 5). b Matrices with full cells filtered to a minimum skeleton length of axon and dendrite of 50 µm each, projecting axon cell types with 50 µm. c Matrices with full cells filtered to a minimum skeleton length of axon and dendrite of 200 µm each, projecting axon cell types with 0 µm. 0 µm refers to axon fragments that only consist of one skeleton node.

*Cell type classification*

| **cell type** | **number of ground truth cells** |
|---|---|
| DA | 26 |
| LMAN | 58 |
| HVC | 53 |
| MSN | 31 |
| STN | 34 |
| TAN | 12 |
| GPe | 13 |
| GPi | 15 |
| LTS | 9 |
| INT1 | 24 |
| INT2 | 16 |
| INT3 | 16 |
| ASTRO | 17 |
| OLIGO | 14 |
| MICRO | 17 |
| MIGR | 14 |
| FRAG | 24 |

Extended Data Table 4: Cell types and their ground truth support for cell classification. This is updated compared to a previous publication [2].

| | redundancy 1 | | redundancy 10 | | redundancy 20 | | redundancy 50 | |
|---|---|---|---|---|---|---|---|---|
| **quantity** | mean | sd | mean | sd | mean | sd | mean | sd |
| **DA** | 0.925 | 0.043 | 0.960 | 0.017 | 0.981 | 0.016 | 0.987 | 0.009 |
| **LMAN** | 0.862 | 0.020 | 0.953 | 0.013 | 0.942 | 0.000 | 0.953 | 0.007 |
| **HVC** | 0.886 | 0.004 | 0.959 | 0.012 | 0.949 | 0.004 | 0.956 | 0.004 |
| **MSN** | 0.884 | 0.006 | 0.995 | 0.007 | 0.995 | 0.007 | 0.995 | 0.007 |
| **STN** | 0.908 | 0.010 | 0.939 | 0.007 | 0.957 | 0.023 | 0.944 | 0.000 |
| **TAN** | 0.823 | 0.036 | 0.973 | 0.019 | 0.973 | 0.019 | 1.0 | 0.000 |
| **GPe** | 0.878 | 0.015 | 0.949 | 0.036 | 0.900 | 0.016 | 0.923 | 0.000 |
| **GPi** | 0.889 | 0.000 | 0.956 | 0.031 | 0.909 | 0.017 | 0.933 | 0.000 |
| **LTS** | 0.522 | 0.046 | 0.850 | 0.035 | 0.85 | 0.035 | 0.825 | 0.035 |
| **INT1** | 0.851 | 0.020 | 0.917 | 0.016 | 0.965 | 0.026 | 0.957 | 0.000 |
| **INT2** | 0.888 | 0.038 | 0.939 | 0.029 | 0.990 | 0.014 | 0.960 | 0.013 |
| **INT3** | 0.938 | 0.045 | 1.0 | 0.000 | 1.0 | 0.000 | 1.0 | 0.000 |
| **ASTRO** | 0.946 | 0.036 | 0.945 | 0.021 | 0.945 | 0.021 | 0.945 | 0.021 |
| **OLIGO** | 0.789 | 0.034 | 0.789 | 0.034 | 0.806 | 0.039 | 0.806 | 0.039 |
| **MICRO** | 0.857 | 0.010 | 0.872 | 0.018 | 0.872 | 0.018 | 0.872 | 0.018 |
| **MIGR** | 0.927 | 0.030 | 0.929 | 0.000 | 0.940 | 0.016 | 0.952 | 0.017 |
| **FRAG** | 0.837 | 0.049 | 0.847 | 0.035 | 0.847 | 0.035 | 0.847 | 0.035 |
| **f1 score macro** | 0.860 | 0.008 | 0.928 | 0.008 | 0.931 | 0.004 | 0.933 | 0.007 |
| **accuracy** | 0.875 | 0.008 | 0.938 | 0.006 | 0.941 | 0.001 | 0.943 | 0.004 |

Extended Data Table 5: Mean and standard deviation of the F1 scores of the 10-fold cross-validation for different cell types and three training repetitions. Redundancy values indicate the number of predictions per neuron. The model used for cell type classification had a redundancy of 20.

| | DA | LMAN | HVC | MSN | STN | TAN | GPe | GPi | LTS | INT1 | INT2 | INT3 | ASTRO | OLIGO | MICRO | MIGR | FRAG |
|---|---|---|---|---|---|---|---|---|---|---|---|---|---|---|---|---|---|
| **DA** | 25.33 | 0 | 0.67 | 0 | 0 | 0 | 0 | 0 | 0 | 0 | 0 | 0 | 0 | 0 | 0 | 0 | 0 |
| **LMAN** | 0 | 57 | 1 | 0 | 0 | 0 | 0 | 0 | 0 | 0 | 0 | 0 | 0 | 0 | 0 | 0 | 0 |
| **HVC** | 0 | 3.33 | 49.67 | 0 | 0 | 0 | 0 | 0 | 0 | 0 | 0 | 0 | 0 | 0 | 0 | 0 | 0 |
| **MSN** | 0 | 0 | 0 | 31 | 0 | 0 | 0 | 0 | 0 | 0 | 0 | 0 | 0 | 0 | 0 | 0 | 0 |
| **STN** | 0 | 0 | 0 | 0 | 33.33 | 0 | 0 | 0 | 0 | 0.67 | 0 | 0 | 0 | 0 | 0 | 0 | 0 |
| **TAN** | 0 | 0 | 0 | 0 | 0 | 12 | 0 | 0 | 0 | 0 | 0 | 0 | 0 | 0 | 0 | 0 | 0 |
| **GPe** | 0 | 0 | 0 | 0 | 0 | 0 | 12 | 1 | 0 | 0 | 0 | 0 | 0 | 0 | 0 | 0 | 0 |
| **GPi** | 0 | 0 | 0 | 0 | 0 | 0 | 1.67 | 13.33 | 0 | 0 | 0 | 0 | 0 | 0 | 0 | 0 | 0 |
| **LTS** | 0 | 0.67 | 0 | 0 | 1 | 0.67 | 0 | 0 | 6.67 | 0 | 0 | 0 | 0 | 0 | 0 | 0 | 0 |
| **INT1** | 0 | 0 | 0 | 0 | 1 | 0 | 0 | 0 | 0 | 23 | 0 | 0 | 0 | 0 | 0 | 0 | 0 |
| **INT2** | 0 | 0 | 0 | 0 | 0 | 0 | 0 | 0 | 0 | 0 | 16 | 0 | 0 | 0 | 0 | 0 | 0 |
| **INT3** | 0 | 0 | 0 | 0 | 0 | 0 | 0 | 0 | 0 | 0 | 0 | 16 | 0 | 0 | 0 | 0 | 0 |
| **ASTRO** | 0 | 0 | 0 | 0 | 0 | 0 | 0 | 0 | 0 | 0 | 0 | 0 | 17 | 0 | 0 | 0 | 0 |
| **OLIGO** | 0 | 0.67 | 0 | 0 | 0.33 | 0 | 0 | 0 | 0 | 0 | 0.33 | 0 | 0 | 9.67 | 2.33 | 0.67 | 0 |
| **MICRO** | 0 | 0 | 0 | 0 | 0 | 0 | 0 | 0 | 0 | 0 | 0 | 0 | 0 | 0 | 17 | 0 | 0 |
| **MIGR** | 0 | 0 | 0 | 0 | 0 | 0 | 0 | 0 | 0 | 0 | 0 | 0 | 0 | 0 | 1 | 13 | 0 |
| **FRAG** | 0.33 | 1.33 | 0.33 | 0.33 | 0 | 0 | 0 | 0 | 0 | 0 | 0 | 0 | 2 | 0.33 | 1.66 | 0 | 17.67 |

Extended Data Table 6: Mean values from the confusion matrix of the 10-fold cross-validation for three different replicates with a redundancy of 20.

*Compartments*

To analyze the cells in detail, the different neuronal compartments were separated in the analysis (see methods). The compartment prediction used was previously described [2]. The prediction is accurate for most cells, however in some cases yielded incorrect compartment associations, like an underprediction of the soma (Extended Data Fig. 13 a-c). When analyzing the surface areas of the somata in all neuronal cell types, the small number of outliers show that extreme cases as the mentioned example are rare (Extended Data Fig. 13 d).

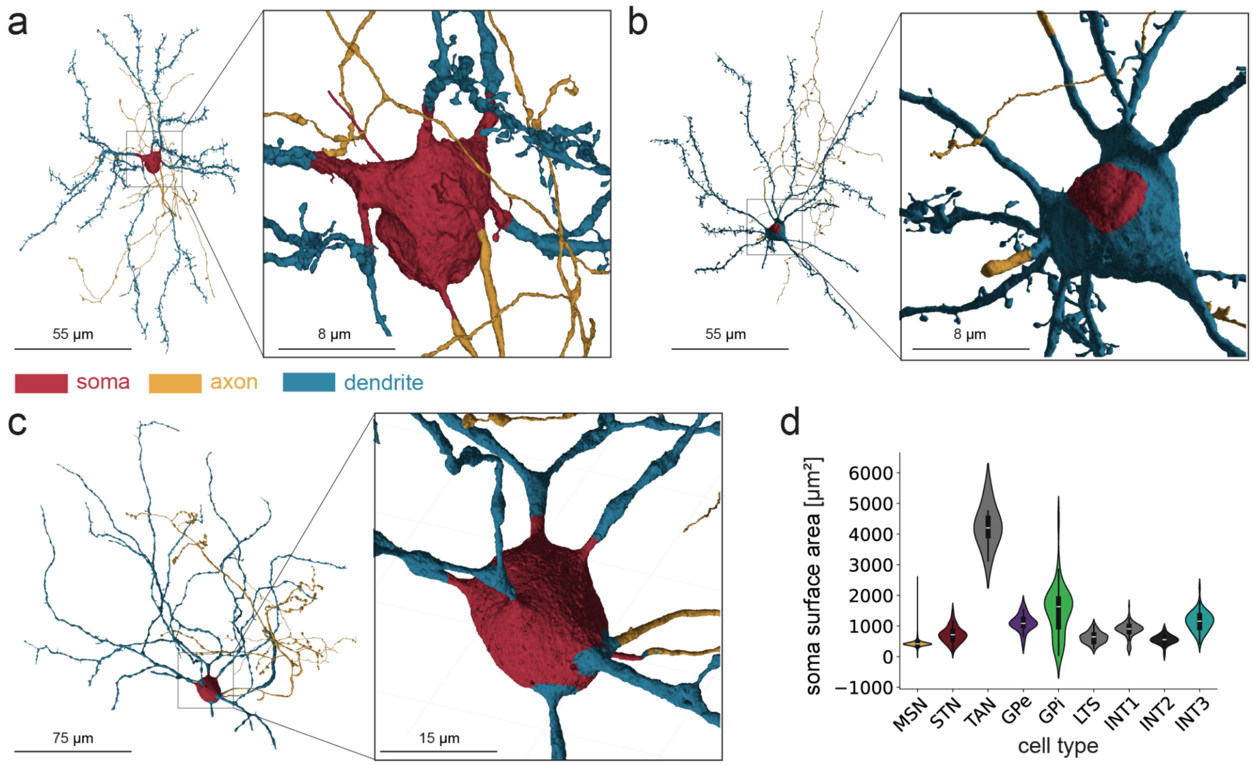


Extended Data Figure 3: Compartment predictions computed previously[2]. a MSN with accurate predictions, soma slightly overpredicted. b MSN with poor soma prediction. c GPi with accurate predictions. d Soma surface area of different cell types.

| cell type | axon pathlength ratio | dendrite pathlength ratio |
|---|---|---|
| **GPe** | 0.49 | 0.49 |
| **GPi** | 0.39 | 0.28 |
| **INT1** | 0.67 | 0.73 |
| **INT2** | 0.22 | 0.57 |
| **INT3** | 0.55 | 0.68 |
| **LTS** | 0.08 | 0.34 |
| **MSN** | 0.77 | 0.73 |
| **STN** | 0.12 | 0.18 |
| **TAN** | 0.04 | 0.23 |

Extended Data Table 7: Ratio of full cells to fragments of a specific compartment per cell type in the dataset. The pathlength ratio calculates the pathlength of that compartment for all full cells versus the summed pathlength of all fragments in that cell type.

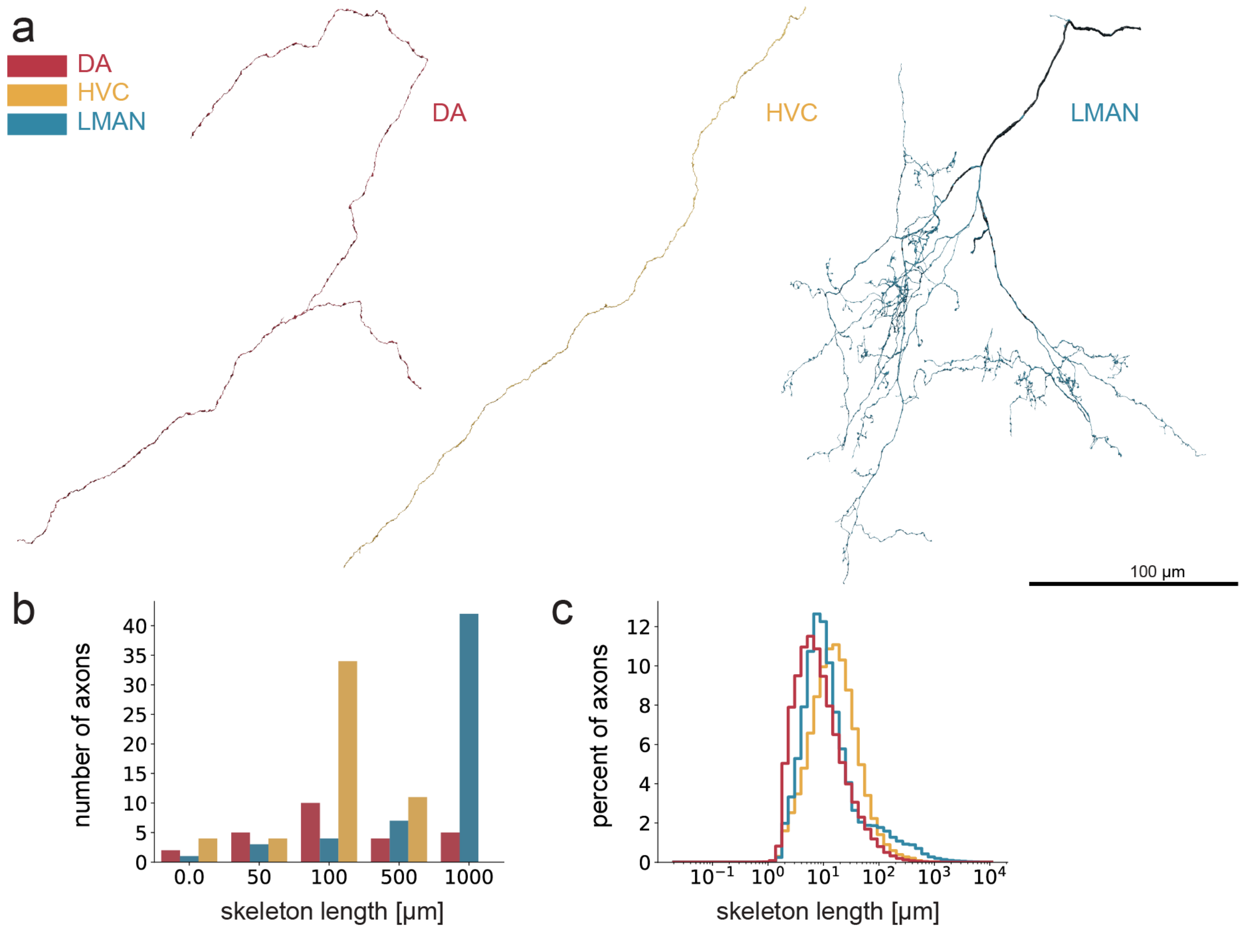


Extended Data Figure 4: Projecting axons to Area X. a Example reconstructions of axons presumably originating from VTA (DA), HVC and LMAN. b Number of axons with their corresponding pathlength in the ground truth. c Number of axons with at least one synapse (size >= 0.1 µm, synapse probability >= 0.6, axo-dendritic)

**Further notes on MSN cells**

In order to see if there are separate MSN populations, different approaches were tested to see if there are subpopulations with different morphologies and if these are related to connectivities.
First, we analyzed whether MSN populations differed in spine density or dendritic length which have been associated with direct and indirect pathway MSNs in mammals[3,4]. There were no clear differences in spine density or dendritic length (Extended Data Fig. 5c) and also no correlation between a preferred connectivity to GPe or GPi and these parameters (Fig. 2d, Extended Data Fig. 5 e). As a control, the same analysis was done with only MSNs whose dendrites were completely inside the dataset (n = 3,052, see methods). There, the correlation of both, the morphological features and the morphology with the connectivity, was weaker ($r = 0.15$ both; $p\text{-value} < 0.001$, Extended Data Fig. 5d, f, g, Extended Data Table 8), and the group only connecting to GPe neurons smaller (24 cells, 0.78%). MSNs only connecting to GPe also had

shorter axons, dendrites and a soma closer to the dataset border compared to MSNs connecting to GPi only or GPe and GPi (p-values < 0.001, except for MSN no GPe vs MSN only GPe: p = 0.027, Extended Data Fig. 5 h-j).

Second, we tested if MSNs could be further subdivided based on a 18-dimensional feature vector which included different morphological parameters and subcellular data such as synapse density, mitochondria density and vesicle cloud density (see methods). The resulting UMAP did not show separate clusters for MSN (Extended Data Fig. 6a), but showed differences in other cell types (Extended Data Fig. 6b) and separated MSNs from the other cell types (Extended Data Fig. 6c). The MSNs in Extended Data Fig. 6a, that were not in the large MSN cloud (first plot, UMAP 1 > 13.5, 45 cells) were manually inspected and two of them (4.4%) were mergers with either another cell or axon but not with glia. The number of mergers in this group is slightly higher than in the full dataset (1.2% out of 500 random cellids, Extended Data Table 3). Their GPe/i specificity was checked and cells belonged to all four groups, with the those projecting to no GPs and only GPe being strongly overrepresented compared to all MSNs (both GPs: 40%, no GPs: 20%, only GPe: 13%, only GPi: 27%). Also in the UMAPs with all cell types (Extended Data Fig. 6c), there is one group separated from the large MSN cloud (second plot, UMAP 2 > 8, 28 cells) which were manually reviewed were all mergers with axon fragments (26) or other cells (2). Their GPe/i specificity was checked and cells belonged to all four groups, but those projecting only to GPe or no GP were overrepresented compared to all MSNs (both GPs: 64%, no GPs: 11%, only GPe: 4%, only GPi: 21%).

When the GP ratio is plotted against the total number of synapses per MSN, the GP ratio approaches 0.55 - 0.6 for extreme cases with MSN > 40 synapses (Extended Data Fig. 3, c, d).

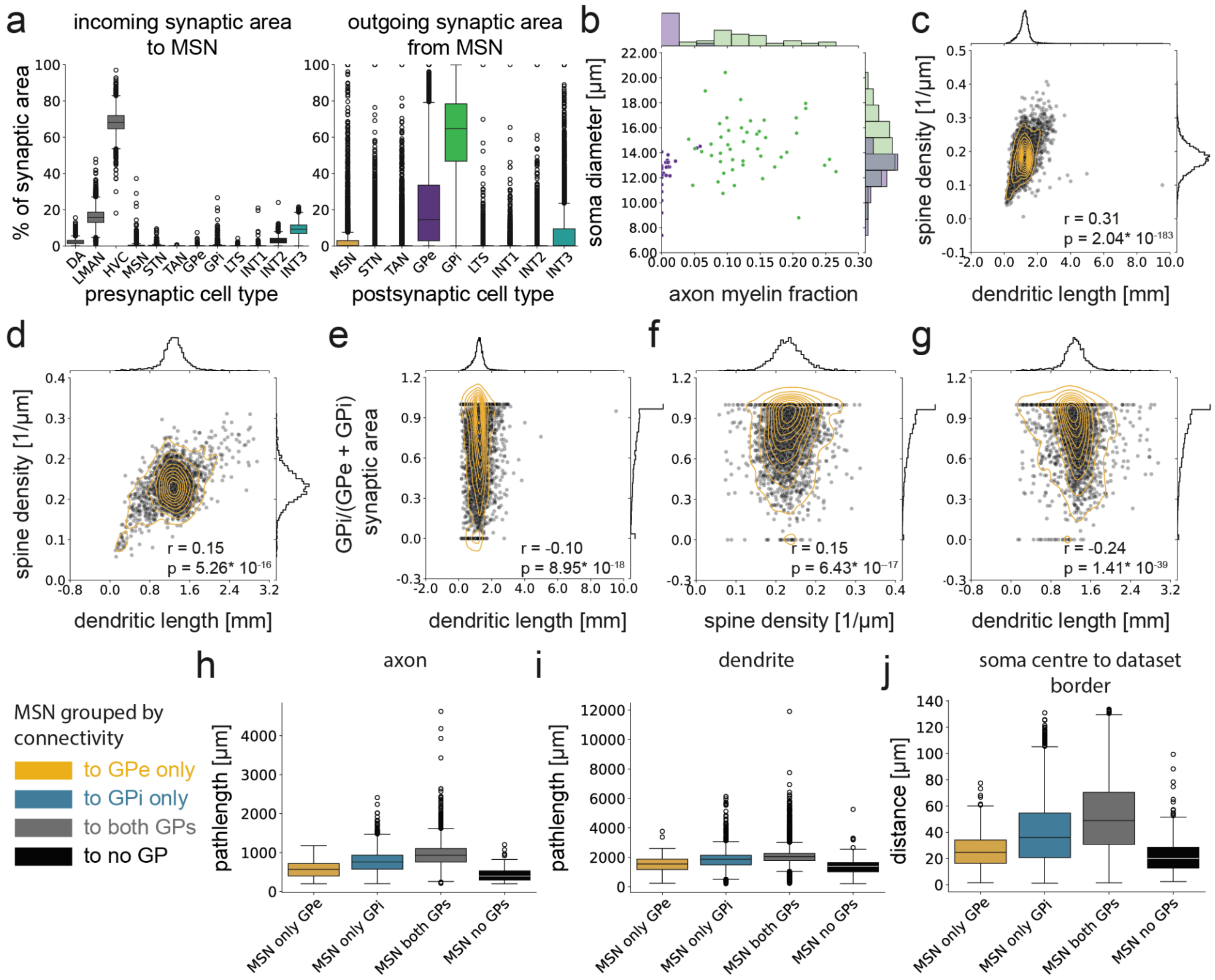


Extended Data Figure 5.a Percentage of incoming and outgoing synaptic area to and from MSN b GPe and GPi separation with the estimated soma diameter and the axon myelin fraction, used for PCA in Fig. 2c. GPe cells are shown in purple, GPi cells in green c There is a broad morphological spectrum of MSNs, with spine density slightly correlated with dendritic length. d Spine density and dendritic length only for MSNs with complete dendrites (n = 3052). e Ratio of GPe and GPi related to dendritic length on all MSN, as in Fig. 2d. f, g Ratio of GPe and GPi related to spine density and dendritic length for MSN without cutoff dendrite, MSN both = 80.3% (2451 cells), MSN GPe only = 0.78% (24 cells), MSN GPi only = 18.2% (556 cells), MSN no GP = 0.69% (21 cells). h Axon skeleton length in MSN grouped by connectivity to GPe and GPi. i Dendritic skeleton pathlength j Distance of the soma center to the nearest dataset border. h-j have significant differences between all groups (Wilcoxon-rank sum test, all $p < 0.005$) except from dendritic length MSN no GP vs MSN GPe only ($p = 0.027$; see Extended Data Table 8).

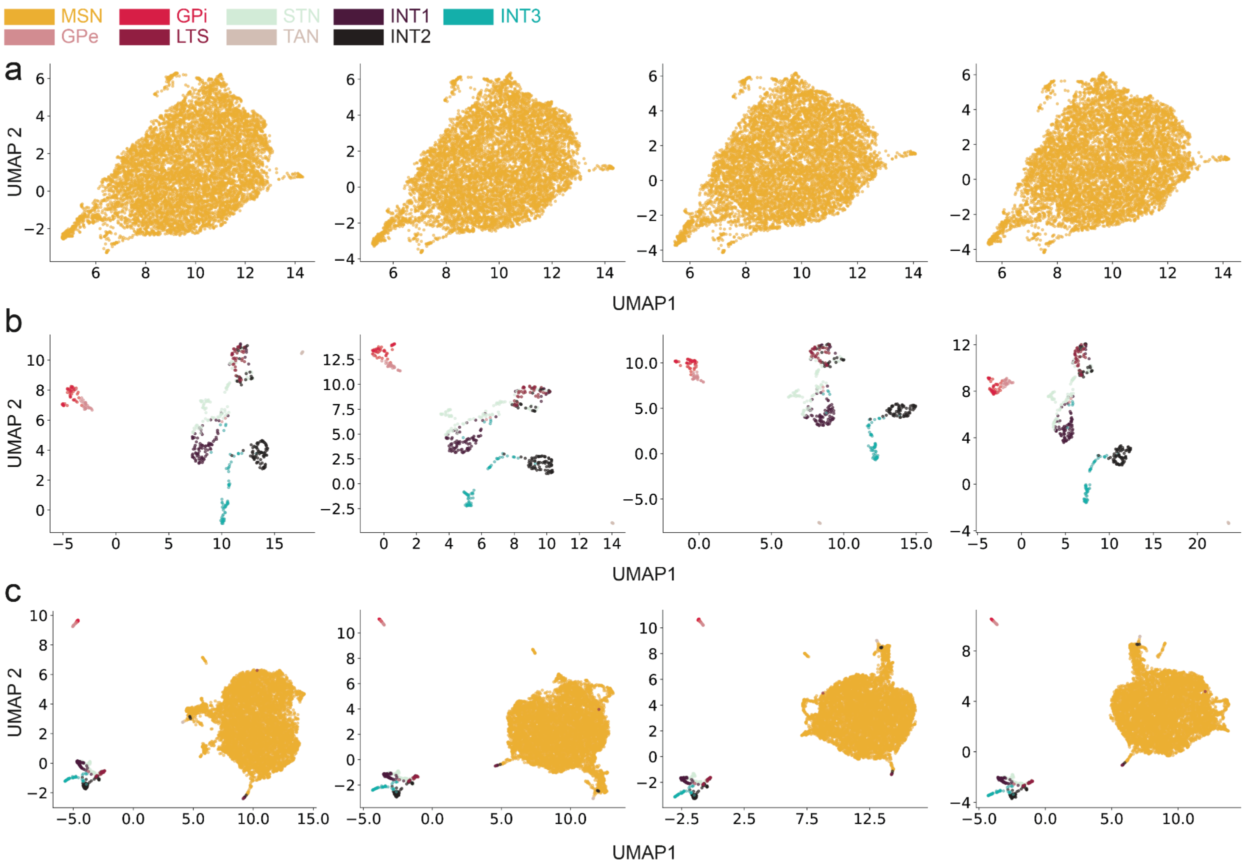


Extended Data Figure 6. UMAP based on an 18 dimensional feature vector based on morphology for MSNs only. b UMAP with the same feature vector as in a, for all full cells. c UMAP with the same feature vector as in d,e with full cells without MSNs.

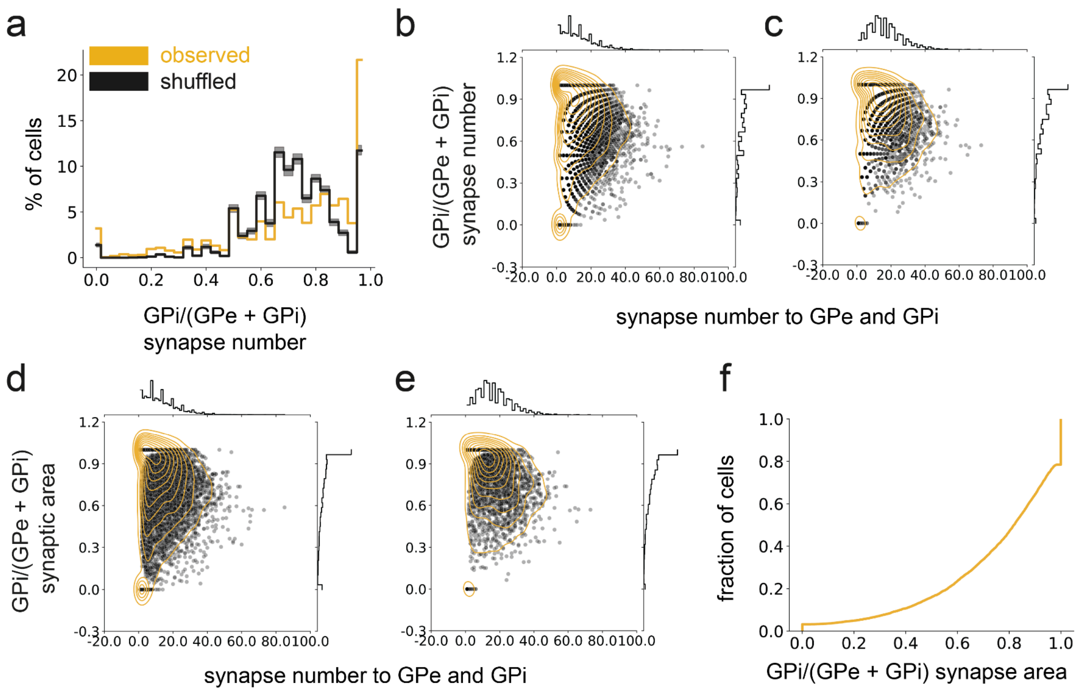


Extended Data Figure 7: a Shuffled vs observed synapse numbers of MSN to GPe and GPi. The shuffled plot represents the mean of 100 iterations, the 95% CI is plotted in gray. b GP ratio of synapse number in relation to the cells total synapse numbers to GPe and GPi for all MSNs. c Same as b for only those with full dendrites. d GP ratio of synaptic area in relation to total synapse number to GPe and GPi for all MSNs. e like d, only full dendrites f Observed synapse area of MSN to GPe and GPi (as in Fig. 2e) plotted as cumulative distribution.

| **compared value** | **statistical test** | **n** | **r (if applicable)** | **p-value** | **figure** |
|---|---|---|---|---|---|
| **MSN all cells, spine density vs dendritic length** | Spearman correlation | 8093 | 0.31 | 2.04*10-183 | Ext Data Fig. 5c |
| **MSN full dendrites only, spine density vs dendritic length** | Spearman correlation | 3052 | 0.15 | 5.26*10-16 | Ext Data Fig.2d |

| | | | | | |
|---|---|---|---|---|---|
| **MSN all cells, GP ratio vs spine density** | Spearman correlation | 7896 | 0.14 | 7.72*10-37 | Fig. 5d |
| **MSN full dendrites only, GP ratio vs spine density** | Spearman correlation | 3031 | 0.15 | 6.43*10-17 | Ext Data Fig 5e |
| **MSN all cells, GP ratio vs dendritic length** | Spearman correlation | 7896 | -0.096 | 8.95*10-18 | Ext Data Fig. 5f |
| **MSN full dendrites only, GP ratio vs dendritic length** | Spearman correlation | 3031 | -0.23 | 1.41*10-39 | Ext Data Fig 5g |
| **MSN groups axon length** | 2-sided Wilcoxon ranksum test, pairwise | only GPe = 253,<br>only GPi = 1700,<br>both = 5943,<br>none = 197 | - | all < 10-14 | Ext Data Fig. 5h |
| **MSN groups dendrite length** | 2-sided Wilcoxon ranksum test, pairwise | | - | all < 10-17 but only GPe vs none = 1.34 * 10-3 | Ext Data Fig. 5i |
| **MSN groups soma centre to dataset border** | 2-sided Wilcoxon ranksum test, pairwise | | - | all < 10-18 but only GPe vs none = 0.027 | Ext Data Fig. 5j |
| **MSN synapse sizes to GPe, to GPi** | 2-sided Wilcoxon ranksum test | MSN - GPe = 28297, MSN - GPi = 76329 | - | 5.3*10-142 | Fig. 2f |
| **MSN number of multi** | 2-sided Wilcoxon ranksum test | MSN - GPe = 343, MSN -GPi = 3516 | - | 1.75*10-121 | Fig. 2g |

| | | | | | |
|---|---|---|---|---|---|
| **synapses to GPe, GPi** | | | | | |
| **MSN summed size per GP cells, to GPe, to GPi** | 2-sided Wilcoxon ranksum test | MSN - GPe = 343, MSN -GPi = 3516 | - | 2.51*10-251 | Fig. 2h |
| **GPe - GPi synapse sizes** | 2-sided Wilcoxon ranksum test | GPe-GPi = 766, GPi - GPe = 124 | - | 0.13 | Fig 3c |
| **GPe - GPi number multi-synaps es** | 2-sided Wilcoxon ranksum test | GPe - GPi = 179, GPi - GPe = 63 | - | 8.47*10-4 | Fig 3d |
| **GPe - GPi summed size per partner cell** | 2-sided Wilcoxon ranksum test | GPe - GPi = 179, GPi - GPe = 63 | - | 0.02 | Fig 3e |
| **GPe-GPe vs GPi - GPi synapse sizes** | 2-sided Wilcoxon ranksum test | GPe = 222, GPi = 6438 | - | 7.69*10-25 | Ext Data Fig. 3a |
| **GPe-GPe vs GPi - GPi number of multi-synaps es** | 2-sided Wilcoxon ranksum test | GPe = 79, GPi = 511 | - | 2.15*10-7 | Ext Data Fig. 3b |
| **GPe-GPe vs GPi - GPi summed syn sizes per partner cell** | 2-sided Wilcoxon ranksum test | GPe = 79, GPi = 511 | - | 3.39*10-14 | Ext Data Fig. 3c |
| **GPe - GPe vs GPi-GPe synapse sizes** | 2-sided Wilcoxon ranksum test | GPe - GPe = 222, GPi -GPe = 124 | - | 9.25*10-6 | Fig. 3c, Ext Data Fig. 8a |
| **GPe - GPe vs GPi-GPe number multi-synaps** | 2-sided Wilcoxon ranksum test | GPe - GPe = 79, GPi - GPe = 63 | - | 0.13 | Fig. 3d, Ext Data Fig. 8b |

| | | | | | |
|---|---|---|---|---|---|
| **es** | | | | | |
| **GPe - GPe vs GPi-GPe summed syn sizes per partner cell** | 2-sided Wilcoxon ranksum test | GPe - GPe = 79, GPi - GPe = 63 | - | 0.43 | Fig. 3e, Ext Data Fig. 8c |
| **GPi - GPi vs GPe - GPi synapse sizes** | 2-sided Wilcoxon ranksum test | GPi - GPi = 6428, GPe - GPi = 766 | - | 0.08 | Fig. 3c, Ext Data Fig. 8a |
| **GPi - GPi vs GPe - GPi number multi-synapses** | 2-sided Wilcoxon ranksum test | GPi - GPi = 511, GPe - GPi = 179 | - | 4.72*10-7 | Fig. 3d, Ext Data Fig. 8b |
| **GPi - GPi vs GPe - GPi summed syn sizes per partner cell** | 2-sided Wilcoxon ranksum test | GPi - GPi = 511, GPe - GPi = 179 | - | 6.37*10-10 | Fig. 3e, Ext Data Fig. 8c |
| **STN - GPe vs STN- GPi synapse sizes** | 2-sided Wilcoxon ranksum test | GPe = 2148, GPi = 7923 | - | 1.32*10-76 | Fig. 4e |
| **STN - GPe vs STN- GPi number multi-synapses** | 2-sided Wilcoxon ranksum test | GPe = 450, GPi = 1066 | - | 7.62*10-17 | Fig. 4g |
| **STN - GPe vs STN- GPi summed syn size per partner cell** | 2-sided Wilcoxon ranksum test | GPe = 450, GPi = 1066 | - | 1.39*10-22 | Fig. 4i |
| **GPe- STN vs GPi - STN synapse size** | 2-sided Wilcoxon ranksum test | GPe = 1969, GPi = 14586 | - | 2.15*10-86 | Fig. 4f |
| **GPe- STN vs GPi - STN number** | 2-sided Wilcoxon ranksum test | GPe = 348, GPi = 1309 | - | 5.65*10-3 | Fig. 4h |

| | | | | | |
|---|---|---|---|---|---|
| **multi-synapses** | | | | | |
| **GPe- STN vs GPi - STN summed syn size per partner cell** | 2-sided Wilcoxon ranksum test | GPe = 348, GPi = 1309 | - | 5.05*10-11 | Fig. 4j |

Extended Data Table 8: Information about the statistical tests performed.

**Further notes on GPe-GPe and GPi-GPi connectivity**

GPe- GPi connectivity is stronger than GPi - GPe. Notably, this innervation pattern arises from three times as many synapses from GPe to GPi (Wilcoxon-ranksum test, $p < 0.001$), while the average synapse size from GPe to GPi is about the same as from GPi to GPe (Wilcoxon-ranksum test, $p = 0.13$, for synapse sizes; $p = 0.02$, for summed synaptic size per cell; Fig. 3c-e).
Synapses from GPi to GPi neurons are larger than in GPe-GPe connections, make more multi-synapses, and have a higher summed synaptic area per cell-pair (Extended Data Fig. 3a-c, Wilcoxon-rank sum test, p-values < 0.005, Extended Data Table 8). Synapses in GPe-GPe connections are smaller than in GPi-GPe connections (Fig 3c, Extended Data Fig. 3a, Wilcoxon-rank sum p-value < 0.005), while they do not differ significantly in the number of multisynaptic connections or in the summed size per cell pair (Fig. 3d, e, Extended Data Fig. 3b, c). GPi-GPi synapses do not differ significantly from GPe-GPi synapses in size (Fig. 3c, Extended Data Fig. 3a), but make more multisynaptic connections and as a result have a higher summed synaptic size per cell pair (Fig. 3d, Extended Data Fig. 3b, c, Wilcoxon-rank sum p-values < 0.005).

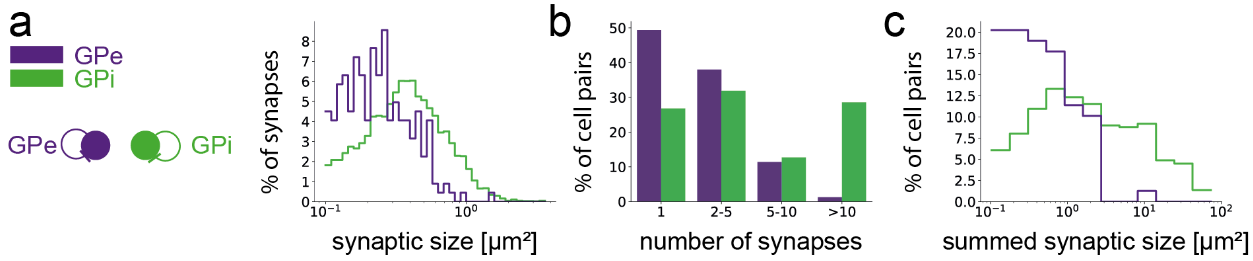


Extended Data Figure 8. Synaptic connectivity within GPe and GPi. a Comparison of synaptic size of individual synapses. b Number of synapses in multisynaptic connections c Summed synaptic area of multisynaptic connections.

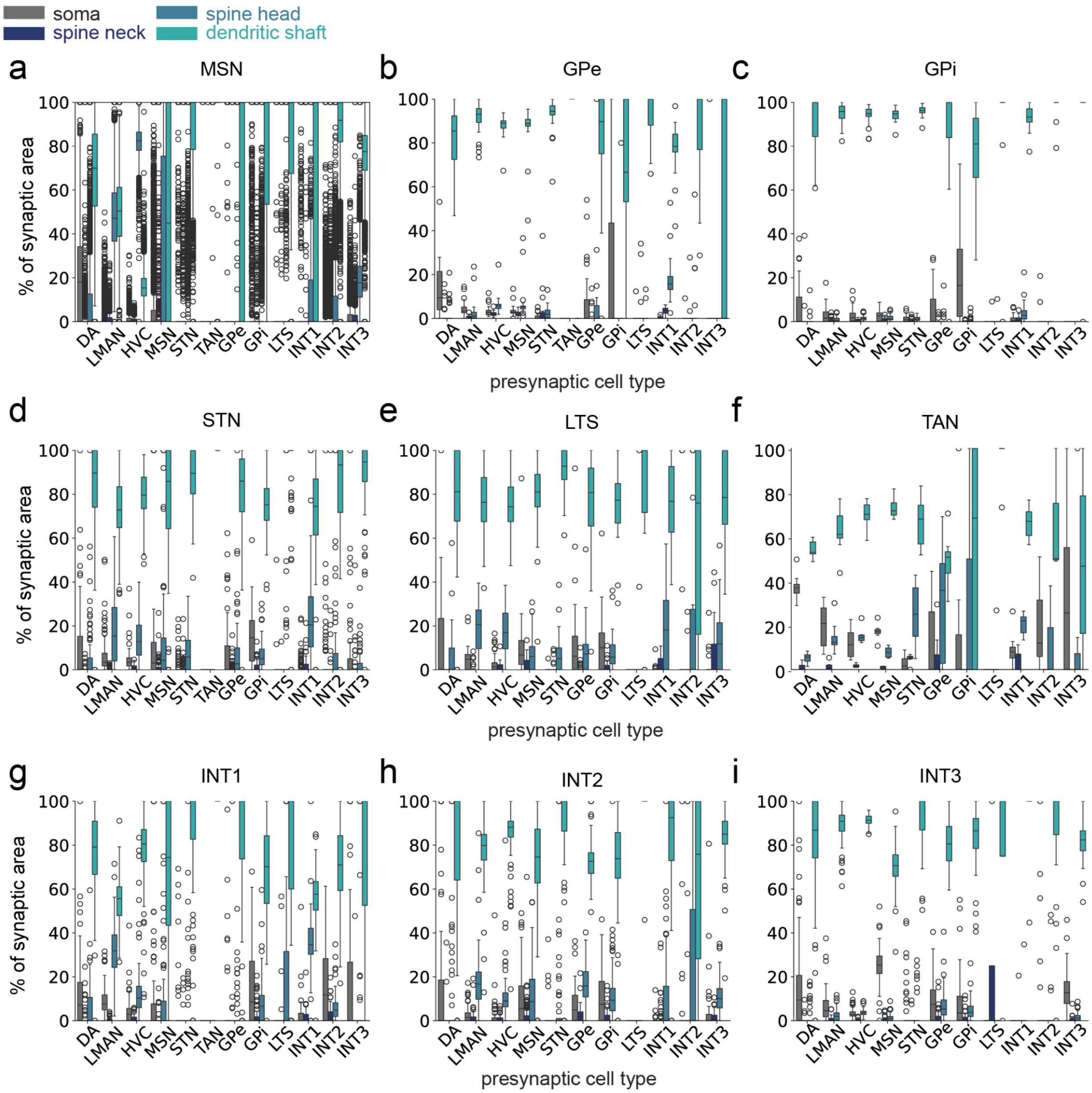


Extended Data Figure 9. Compartment specific connectivity for all the neuron types in the dataset, that reside in Area X. Plots show the percentage of synaptic area per presynaptic cell type, so compartments for each presynaptic cell type add up to 100%. Each data point is a postsynaptic cell.a-i Compartment specific connectivity for different postsynaptic cell types. In c, e, f, h, i, TAN is missing because no TAN-TAN synapses were detected ($n_{TAN}$ = 8).

### Further notes on reciprocity of STN, GPe and GPi

Since 18% of STN cells do not project to either GPe or GPi (see Extended Data Table 1), the connectivity of these cells was analyzed. Both the fraction of incoming and the fraction of outgoing synaptic area to and from the MSNs is higher compared to all STNs (incoming: 19.8 +-

12.9% vs all STN: 7.4+- 10.8%; outgoing: 32.9 +- 39.6% vs all STNs: 19.1 +- 32.2%, Fig. 4 d, Extended Data Fig. 4a). STN makes recurrent connections to both GPe and GPi, to analyze if this is a cell-specific recurrency two parameters were looked at, the strongest partner reciprocity and the overlap in synaptic area. Strongest partner reciprocity was set to true only if the strongest input (largest summed synaptic area) of a cell was also the strongest output. Synaptic area overlap was calculated as the fraction of the summed synaptic area of those cells, that were both on the input and output side, divided by the total area of input or output that cell received from the other cell type. For comparison, MSN and TAN cells were also analyzed. Note that all complete GPe, GPi and TAN cells (Extended Data Table 2) were included in the analysis, while only a small fraction of MSNs were (<0.5%). Both the strongest partner reciprocity (Extended Data Fig. 4b) and the overlap fraction of synaptic areas (Extended Data Fig. 4c-h) show that while STN-GPi and STN-GPe connections result in cell-specific recurrency, this is not the case for MSN-TAN connections. This could also be an effect of cell size, as MSNs are much smaller than all the other cell types analyzed. As STN makes recurrent connections with GPe and GPi, these were compared on the level of individual cells and synapses. In addition to differences between synapses to and from GPe/i, we also investigated if this recurrency exists on a single cell level, i.e. whether individual neuron pairs form feedback loops, or whether connectivity in one direction suppresses it backwards. The synaptic area connections to and from the same cells also make up the majority of in-and outgoing cells, for all tested combinations except GPe outgoing synapses to STN (STN - GPi: STN in 90%; out 62% GPi in 75%, out 75; STN-GPe: STN in 76%, out 60% GPe in 73%; out 48%, Extended Data Fig. 10c-f). While for STN-GPi the majority of cells follows this distribution, for STN-GPe a quarter of STN cells have very little overlap (see Extended Data Fig. 10c-f). This indicates that the recurrent connectivity identified on the cell type level can also be found to some degree on the level of individual neurons, predicting the existence of functional excitatory - inhibitory feedback loops between STN and GPi neurons. For the STN-GPe connections, 18.6% of STN neurons and 18.5% of GPe neurons have the same neuron as their strongest input and output partner. For STN-GPi connections, this is 14.1% and 19.1% respectively.

| **STN connectivity** | **number of STN cells** | **% of STN cells** |
|---|---|---|
| **GPe and GPi** | 58 | 56.86 |
| **no GP** | 18 | 17.65 |
| **STN only GPe** | 18 | 17.65 |
| **STN only GPi** | 8 | 7.84 |

Extended Data Table 9: Number of STN cells projecting to cells of classes GPe and GPi, only one of them, or none.

| connection | cell number input and output | fraction cells strongest partner reciprocity | median number of partner cells incoming | median number of partner cells outgoing | median fraction synapse sum size overlap incoming | median fraction synapse sum size overlap outgoing |
|---|---|---|---|---|---|---|
| **STN - GPi** | 64, 47 | 0.141, 0.191 | 13, 28 | 21, 23 | 0.90, 0.75 | 0.62, 0.75 |
| **STN - GPe** | 70, 27 | 0.186, 0.185 | 5, 13 | 4, 16 | 0.76, 0.73 | 0.60, 0.48 |
| **MSN - TAN** | 29, 8 | 0.276, 0 | 1, 236.5 | 1, 18.5 | 0.30, 0.0029 | 0.27, 0.063 |

Extended Data Table 10: Cell-specific recurrency for connections between different cell types. The number of input and output cells refers to the number of cells that both project to and receive input from the other cell type. Note that all full cells of GPe, GPi and TAN do this for the given projections, while only a small fraction of MSNs (<0.5%) project to TAN and receive input from them. Strongest partner reciprocity refers to the number of cells where the strongest input is also the strongest output. The fraction of synaptic summed size overlap was calculated as the summed area of synapses that are from cells that are both inputs and outputs, divided by either the sum of the summed area of all inputs from that cell type (with cells in the analyses) or all outputs.

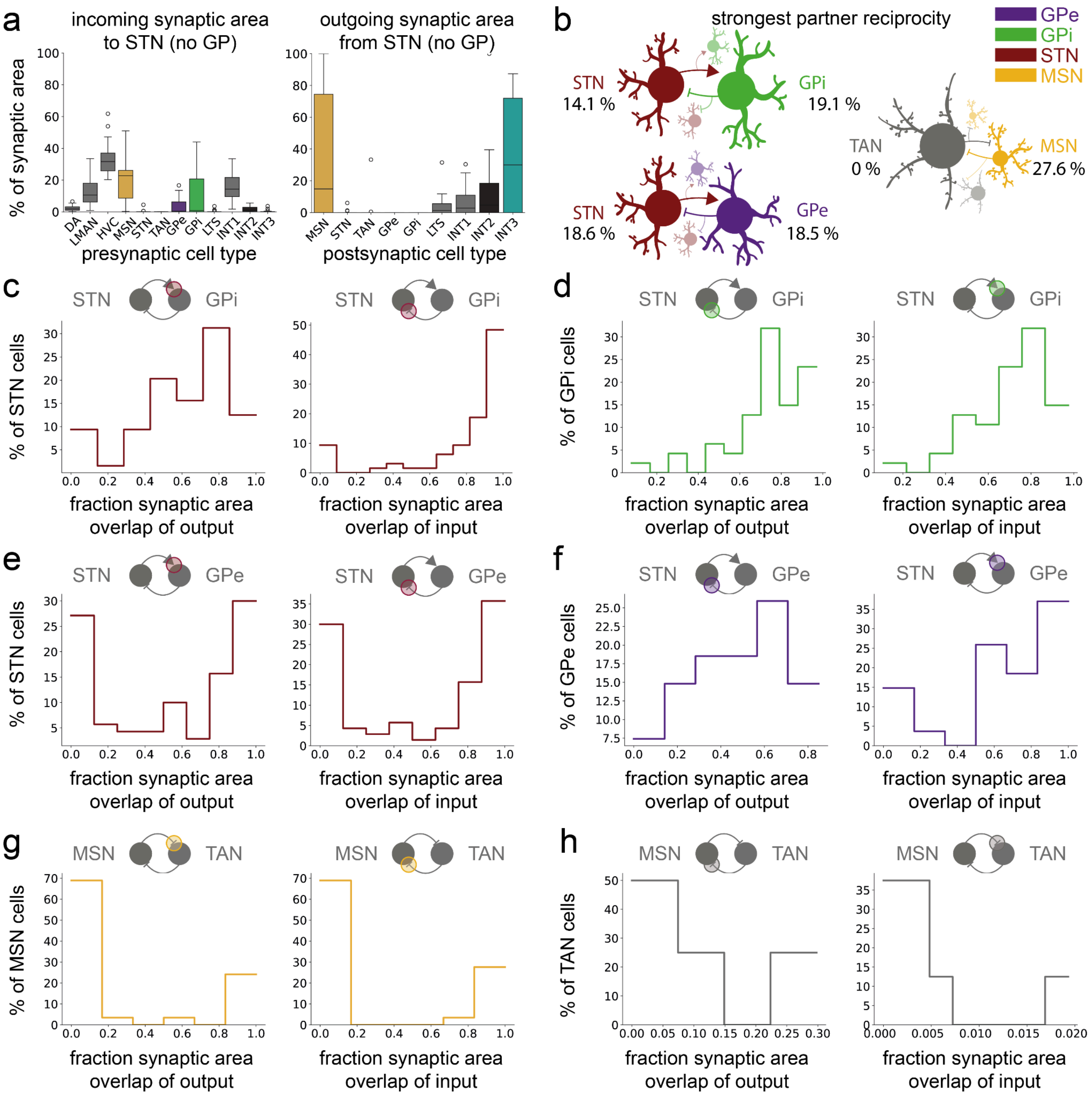


Extended Data Figure 10: STN connectivity and recurrency. a Incoming and outgoing synaptic area of STN cells that do not project to GPe or GPi (18%, see Extended Data Table 1). b Strongest partner reciprocity: fraction of cells whose largest synaptic input area comes from the cell to which their largest synaptic output area goes in the specific connections.c -h Fraction of synaptic area of cells that are both inputs and outputs compared to the total summed synaptic area for all cells that are potential targets for recurrent connections, either incoming or outgoing. The results of b-h are also summarized in Extended Data Table 2.

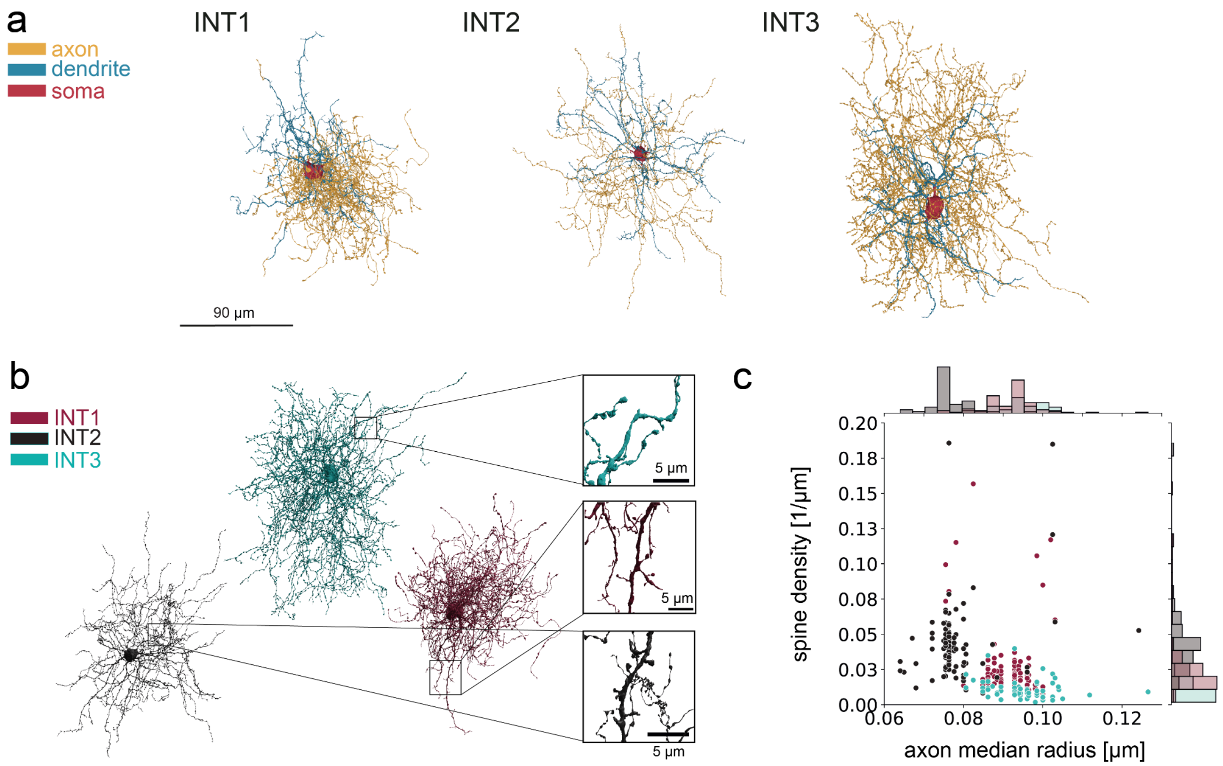


Extended Data Figure 11. a Renderings of INT1, INT2 and INT3 with different colors for axon, dendrite and soma. The cells are the same as in Fig. 5a. b Cells from **a)** with zoom on their spine density. c Spine density and axon median radius for the three cell types, two out of the four parameters used for the PCA in Fig. 5c, soma diameter and axonal mitochondria volume density are shown in Fig. 5b.

| **cell type** | **number of manually inspected cells** | **cells with min. 1 autapse** | **percentage of cells with autapses** |
|---|---|---|---|
| **GPe** | 26 | 8 | 31% |
| **GPi** | 42 | 17 | 40% |
| **INT1** | 88 | 9 | 10% |
| **INT2** | 96 | 2 | 2% |
| **INT3** | 63 | 34 | 54% |
| **LTS** | 36 | 0 | 0% |
| **MSN** | 50 | 0 | 0% |
| **STN** | 97 | 0 | 0% |
| **TAN** | 7 | 0 | 0% |

Extended Data Table 11: Fraction of cells with a manually verified autapse in different cell types. For GPe, GPi, LTS, STN, TAN, INT1, INT2 and INT3, all cells used for analysis were inspected that had complete soma within the data. For MSNs, a random subset was inspected.

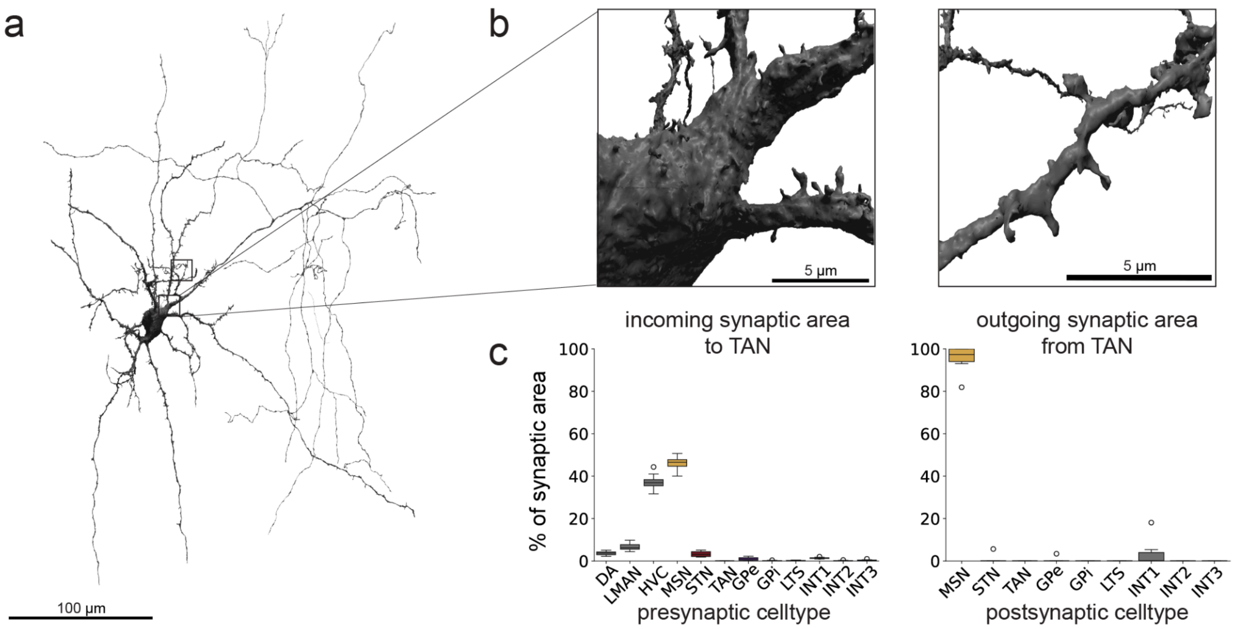


Extended Data Figure 12. a Rendering of a TAN cell. b Spines on the soma and dendrite of a TAN cell. c Connectomic characterization of their synaptic inputs and outputs as percentage of synaptic area per cell type.

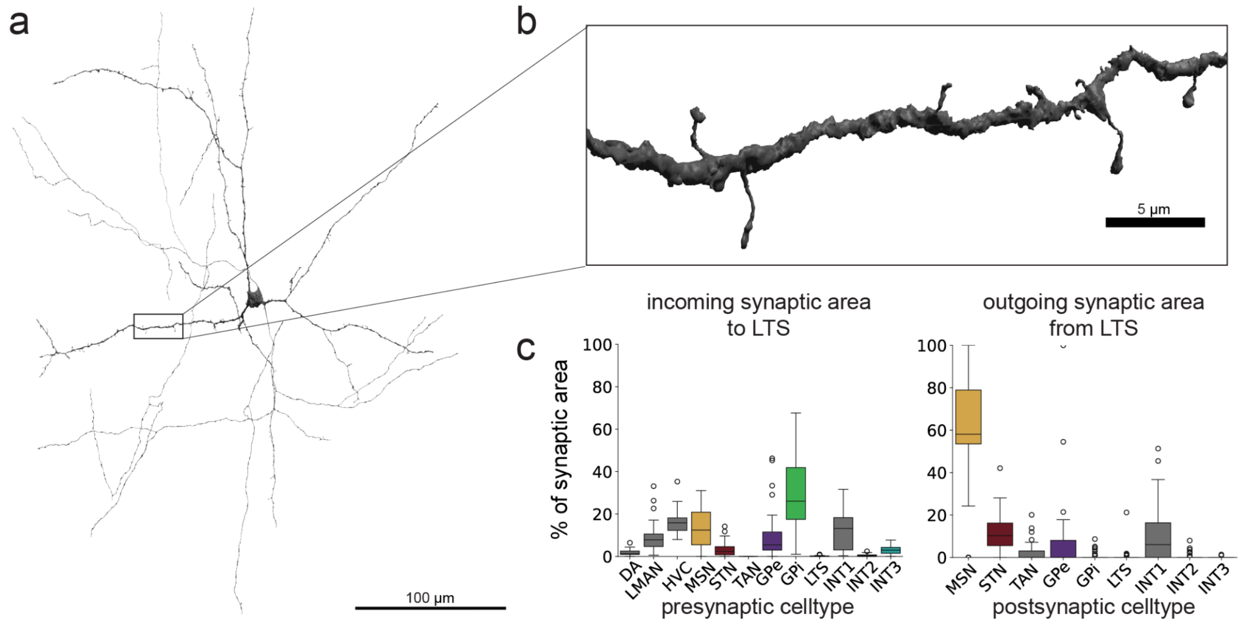


Extended Data Figure 13. a Rendering of an LTS cell. b Sparsely spiny dendrites of the LTS cell. c Connectomic characterization of their synaptic inputs and outputs as percentage of synaptic area per cell type.

**Further notes on LTS cells:**

The LTS cells have a low classical synapse density (data not shown) and little synaptic influence, as measured by us, on all other cell types (Fig. 6c). This is consistent with findings in the mammalian striatum[5,6]. While LTS in the mammalian striatum also mainly project to MSN and receive cortical input, our results differ with GPi as their major input source (Extended Data Fig. 13). Our results suggest a potential feedback and feedforward role.

A modulatory role with potential volume release, similar to dopaminergic axons and TANs, has been suggested for LTS, in this case with a potential release of somatostatin and neuropeptide Y[5,6]. In our data, as is the case for putative DA axons and TANs, a large number of axonal vesicles of LTS axons can be found outside of clear synaptic loci (data not shown).

**Supplementary References**

1. Dorkenwald, S. *et al.* Automated synaptic connectivity inference for volume electron microscopy. *Nat. Methods* **14**, 435–442 (2017).

2. Schubert, P. J. *et al.* SyConn2: dense synaptic connectivity inference for volume electron microscopy. *Nat. Methods* **19**, 1367–1370 (2022).

3. Gagnon, D. *et al.* Striatal Neurons Expressing D and D Receptors are Morphologically Distinct and Differently Affected by Dopamine Denervation in Mice. *Sci. Rep.* **7**, 41432 (2017).

4. Gertler, T. S., Chan, C. S. & Surmeier, D. J. Dichotomous anatomical properties of adult striatal medium spiny neurons. *J. Neurosci.* **28**, 10814–10824 (2008).

5. Gittis, A. H., Nelson, A. B., Thwin, M. T., Palop, J. J. & Kreitzer, A. C. Distinct roles of GABAergic interneurons in the regulation of striatal output pathways. *J. Neurosci.* **30**, 2223–2234 (2010).

6. Ibáñez-Sandoval, O. *et al.* A novel functionally distinct subtype of striatal neuropeptide Y interneuron. *J Neurosci* **31**, 16757–16769 (2011).
